# Integrative proteomics reveals exceptional diversity and versatility of mammalian humoral immunity

**DOI:** 10.1101/2020.08.21.261917

**Authors:** Yufei Xiang, Zhe Sang, Lirane Bitton, Jianquan Xu, Yang Liu, Dina Schneidman-Duhovny, Yi Shi

**Author notes:** Equal contribution (Yufei Xiang, Zhe Sang).

## Abstract

The humoral immune response is essential for the survival of mammals. However, we still lack a systematic understanding of the specific serologic antibody repertoire in response to an antigen. We developed a proteomic strategy to survey, at an unprecedented scale, the landscapes of antigen-engaged, serum-circulating repertoires of camelid heavy-chain antibodies (hcAbs). The sensitivity and robustness of this technology were validated using three antigens spanning orders of magnitude in immune response; thousands of divergent, high-affinity hcAb families were confidently identified and quantified. Using high-throughput structural modeling, cross-linking mass spectrometry, mutagenesis, and deep learning, we mapped and analyzed the epitopes of > 100,000 antigen-antibody complexes. Our results revealed a surprising diversity of high-affinity hcAbs for specific antigen binding on a variety of dominant epitopes. hcAbs perfect both shape and charge complementarity to target challenging antigens specifically; they can rapidly evolve to recognize a conserved, promiscuous cellular protein interaction interface, unraveling the convergent force that drives protein-protein interactions.

## Introduction

In response to an antigenic challenge, mammals produce serologic antibodies with outstanding affinity and selectivity towards the antigen (Chaplin, 2010). The humoral immune response is universal and critical for all mammals, including humans, for surviving countless pathogenic challenges. Despite enormous efforts in characterizing specific binary antigen-antibody interactions (Inbar et al., 1972; Sela-Culang et al., 2013), we still lack a comprehensive picture of the circulating antibody repertoire for antigen binding. The complexity and diversity of such a collection, the epitopes it targets, and the mechanisms underlying high-affinity binding, and the nature of antigenicity remain fully explored. Systematic, large-scale identification and characterization of antigen-engaged antibody proteomes, including high-throughput structure determination of antigen-antibody complexes, while still beyond the reach of current technologies (Cheung et al., 2012; Egloff et al., 2019; Fridy et al., 2014; McMahon et al., 2018; Wine et al., 2013), will provide novel insights into mammalian humoral immunity and the disease mechanisms. The development of cutting-edge tools that facilitate high-quality antibody analysis may also help advance novel antibody-based therapeutics and disease diagnostics (Baran et al., 2017 35; Chevalier et al., 2017; Sircar et al., 2011).

Camelid heavy-chain-only antibodies (hcAbs) and their natural antigen-binding fragments (VHHs or nanobodies) have recently emerged as a new class of antibodies for biomedical applications (Muyldermans, 2013). Nanobodies (Nbs) form conserved, soluble, and compact core structures that are composed of four framework regions (FRs). Complementarity-determining regions (i.e., CDR1, CDR2, and CDR3) form unique hypervariable loops supported by the robust β-sandwich fold and are essential for antigen interaction (Desmyter et al., 1996; Hamers-Casterman et al., 1993). The small size, high solubility, stability, and structural robustness of Nbs greatly facilitate high-throughput microbial production, biophysical and structural characterization, making them an excellent model system to investigating mammalian humoral immunity (Beghein and Gettemans, 2017).

Here we developed a pipeline for large-scale identification, quantification, classification, and high-throughput structural characterization of antigen-specific Nb repertoires. The sensitivity and robustness of this approach were validated using antigens that span three orders of magnitude in immune response including a small, weakly immunogenic antigen derived from the mitochondrial membrane. Tens of thousands of diverse, specific Nb families were confidently identified and quantified according to their physicochemical properties; a significant fraction had a sub-nM affinity. Using high-throughput structural modeling, structural proteomics, and deep learning, we have surveyed the structural landscapes of >100,000 antigen-Nb complexes to advance our understanding of the humoral immune response. Our big data has revealed a surprising efficiency, specificity, diversity, and versatility of the mammalian humoral immunity.

## Results

### The superiority of chymotrypsin for Nb proteomics analysis

We amplified the variable domains of HcAb (V_H_H/Nb) cDNA libraries from the B lymphocytes of two *lama glamas*, recovering 13.6 million unique Nb sequences in the databases by the next-generation genomic sequencing (NGS) (DeKosky et al., 2013). Approximately half a million Nb sequences were aligned to generate the sequence logo (**Fig 1A**). CDR3 loops have the largest sequence diversity and length variation providing excellent specificity for Nb identifications (**Fig 1B-C**). *In silico* analysis of Nb databases revealed that trypsin predominantly produced large CDR3 peptides due to the limited number of trypsin cleavage sites on Nbs. Unlike human or murine IgGs, the presence of proline (**Fig 1A**, position 84) C terminal to lysine (position 83) on the FR3 region appears to be unique to camelid HcAbs/Nbs, rendering efficient trypsin cleavage difficult (**Fig S1A**). The majority of the resulting CDR3 residues (77%) were covered by large tryptic peptides of more than 2.5 kDa (**Fig 1D-E**), which are suboptimal for proteomic analysis (**Fig S1B**). In comparison, chymotrypsin, which is infrequently used for proteomics cleaving specific aromatic and hydrophobic residues, appears to be more suitable (**Methods, Fig 1A**, **S1C**). 91% of CDR3 sequences can be covered by chymotryptic peptides less than 2.5 kDa (**Fig 1D-E**). Random selection and simulation confirmed that significantly more CDR3 sequences could be covered by chymotrypsin than trypsin (**Fig 1F**). Moreover, there was a small overlap (~9%) between the two enzymes, indicating their good complementarity for efficient Nb analysis.

**Fig 1.**
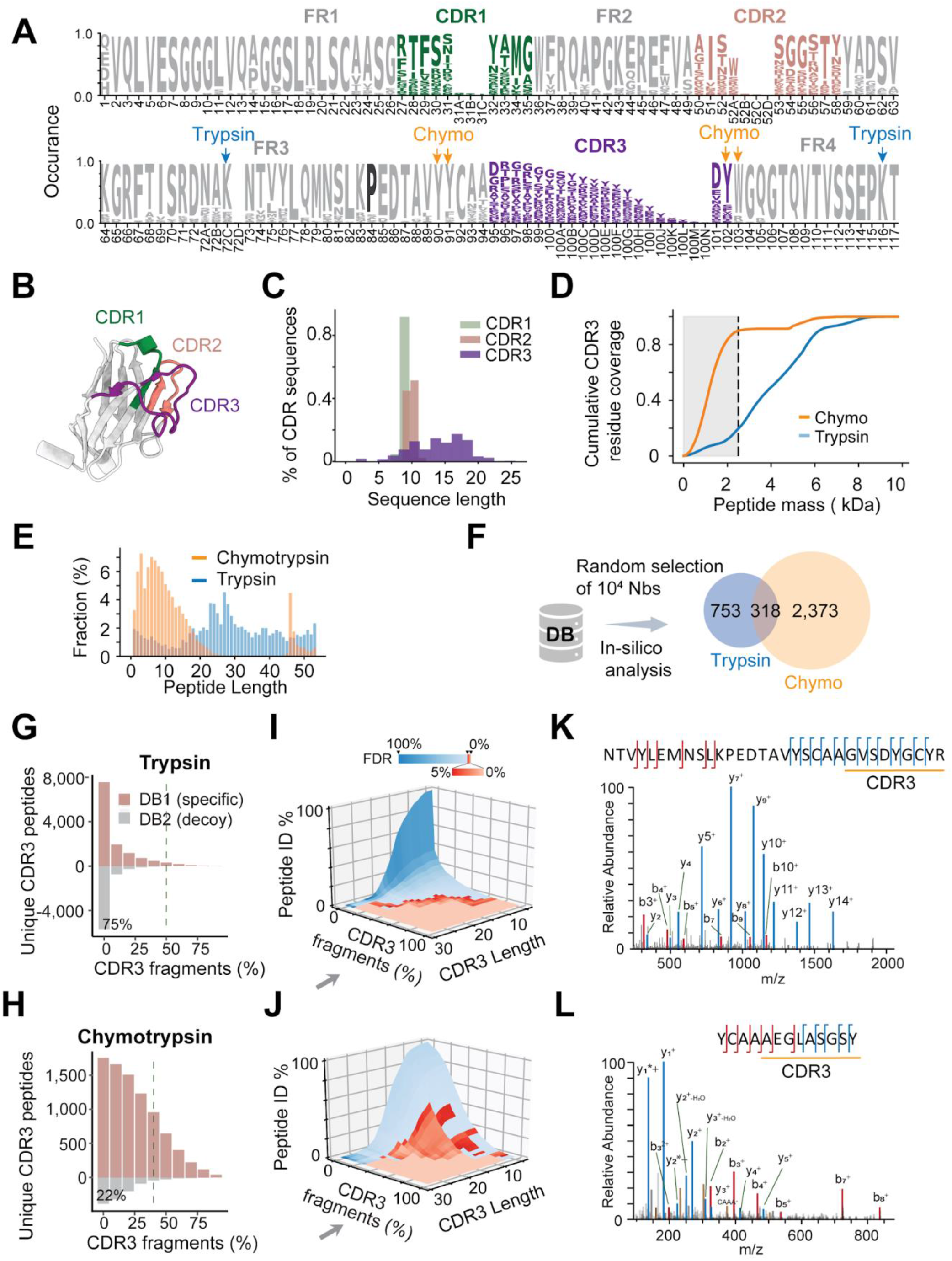
*In silico* analysis of a NGS Nb database reveals the superiority of chymotrypsin for Nb proteomics. A) Logo display of amino acid occurrence of approximately 0.5 million Nb sequences from the NGS database. CDR: complementarity-determining region. FR: framework region. Arrows in blue and orange denote the abundant cleavage sites by trypsin and chymotrypsin, respectively. Proline at position 84 is bolded. B) A Nb crystal structure (PDB: 4QGY). CDR loops are color coded. C) The length distributions of CDRs of the database. D) *In silico* digestion of the Nb database by two proteases and a cumulative plot of corresponding peptide masses. E) The length distributions for both trypsin and chymotrypsin digested CDR3 peptides. F) Complementarity of trypsin and chymotrypsin for Nb mapping based on simulation. 10,000 Nbs with unique CDR3 sequences were randomly selected and *in silico* digested to produce CDR3 peptides. Peptides with molecular weights of 0.8- 3 kDa and with sufficient CDR3 coverage (≥ 30%) were used for Nb mapping. G-H) Evaluations of unique CDR3 peptide identifications (G: trypsin; H: chymotrypsin) based on the percentage of CDR3 fragment ions that were matched in the MS/MS spectra. CDR3 peptides were identified by database search using either the “target” database (in salmon) or the “decoy” database (in grey). I-J) 3D plots of the normalized CDR3 peptide identifications from the target database search, the percentages of CDR3 fragmentations, and CDR3 length. FDR: false discovery rate. FDRs of CDR3 identifications are colored on the 3D plots. The color bar shows the scale of FDR. FDR below 5% are presented in gradient red. I: analysis by trypsin; J: analysis by chymotrypsin. K-L) Representative high-quality MS/MS spectra of CDR3 peptides by trypsin and chymotrypsin.

We hypothesize that the estimated false discovery rate (FDR) of CDR3 identifications could be inflated due to the large database size and the unusual Nb sequence structure. To test this, we proteolyzed antigen-specific HcAbs with trypsin or chymotrypsin. We employed a state-of-the-art search engine Proteome Discovery (Sequest HT) for identification using two different databases: a specific “target” database derived from the immunized llama, and a “decoy” database of similar size from an irrelevant llama with literally no identical sequences (**Fig S1D**)Thus, any CDR3 peptides identified from the decoy database search were considered as false positives (Elias and Gygi, 2007). A surprisingly large number of false-positive CDR3 peptides were nonspecifically identified from the decoy database search. We found that these spurious peptide-spectrum-matches generally contained poor MS/MS fragmentations on the CDR3 fingerprint sequences (**Fig S1E-F**). The vast majority (95%) of these erroneous matches could be removed by using a simple fragmentation filter that we have implemented, requiring a minimum coverage of 50% (by trypsin, **Fig 1G**) and 40% (by chymotrypsin, **Fig 1H**) of the CDR3 high-resolution diagnostic ions in the MS2 spectra (**Fig 1K-L**). The filter was further optimized based on the CDR3 length (**Fig 1I-J**) before integrating into an open-source software Augur Llama (**Fig S2A-C**) that we developed for reliable Nb proteomic analysis.

### Development of an integrative proteomics pipeline for Nb discovery and characterization

We developed a platform for comprehensive quantitative Nb proteomics and high-throughput structural characterizations of antigen-Nb complexes (**Fig 2A**). A domestic camelid was immunized with the antigens of interest. The Nb cDNA library was then prepared from the blood and bone marrow of the immunized camelid (Fridy et al., 2014). NGS was performed to create a rich database of >10^7^ unique Nb protein sequences (**Fig S2D-E**). Meanwhile, antigen-specific Nbs were isolated from the sera and eluted using step-wise gradients of salts or pH buffers. Fractionated HcAbs were efficiently digested with trypsin or chymotrypsin to release Nb CDR peptides for identification and quantification by nanoflow liquid chromatography coupled to high-resolution MS. Initial candidates that pass database searches were annotated for CDR identifications. CDR3 fingerprints were filtered to remove false positives, their abundances from different biochemical fractions were quantified to infer the Nb affinities, and assembled into Nb proteins – all of the above steps were automated by Augur Llama. In parallel, high-throughput structural docking (Schneidman-Duhovny et al., 2005), cross-linking mass spectrometry (CXMS) (Chait et al., 2016; Leitner et al., 2016; Rout and Sali, 2019; Yu and Huang, 2018), and mutagenesis were integrated into our pipeline for accurate epitope mapping and verification. A deep-learning model was developed to learn the features associated with the high-affinity antibodies (**Methods**).

**Fig 2.**
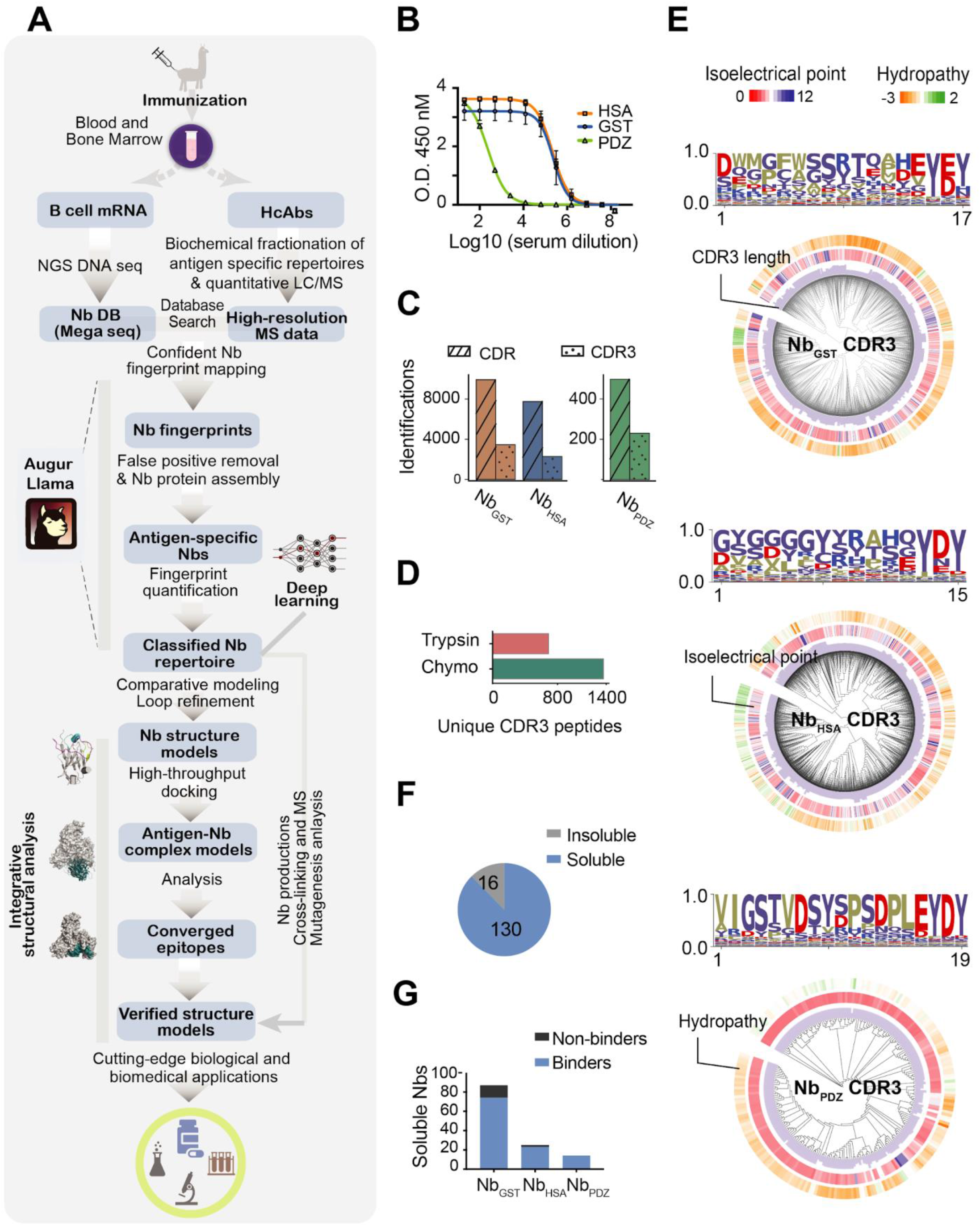
Schematics of a robust pipeline for analysis of antigen-engaged Nb proteomes. A) Schematics of the pipeline for Nb proteomics. The pipeline consists of three main components: camelid immunization and purification of antigen-specific Nbs, proteomic analysis of Nbs (facilitated by a software Augur Llama and deep-learning), and high-throughput integrative structural analysis of antigen-Nb complexes. B) ELISA analysis of the camelid immune responses of three different antigens. C) Identifications of unique CDR combinations and unique CDR3 sequences for different antigens. D) A comparison between trypsin and chymotrypsin for CDR3 mapping of Nb_GST_. E) Phylogenetic analysis and logo plots of CDR3 sequences from three antigen-specific repertoires. CDR3 length is shown in bar plot, and the pI and hydropathy are shown in the heatmaps. Hydrophobic, negatively charged, positively charged, and hydrophilic or neutral residues are shown in olive, red, blue, and indigo, respectively. F) The solubility of the randomly selected antigen-specific Nbs. G) Verifications of the selected Nbs for antigen binding.

### Robust, in-depth, and high-quality identifications of antigen-specific Nbs

To validate this technology and to improve our understanding of the humoral immune response, we immunized llamas with three antigens that elicit several orders of magnitude in immunogenicity (**Fig 2B**): human serum albumin (HSA) (Curry et al., 1998), glutathione S-transferase (GST) (Sheehan et al., 2001), both of which have rich geometric surface features; and a small, conserved, weakly immunogenic PDZ domain derived from the mitochondrial outer membrane protein (OMP25) (Doyle et al., 1996).

A surprisingly large number of 5,969 unique CDR3 families (3,453, 2,286, and 230 individual CDR3 families for GST, HSA, and PDZ, respectively) corresponding to 18,159 unique CDR combinations of serologic Nbs were discovered using our approach (**Fig 2C, S2H**). Each Nb repertoire was antigen-specific and had remarkable diversity in terms of sequence length, hydrophobicity, and isoelectric point (pI) of the CDR3 fingerprints (**Fig 2E**). We confirmed that chymotrypsin provided the most useful fingerprint information for Nb identification from the various proteases tested (**Fig 2D**). A random set of 146 Nbs with unique CDR3s was selected for production in *E.coli*, of which 89% were highly soluble (**Fig 2F-G**), and a fraction was analyzed for antigen binding by complementary assays including immunoprecipitation, ELISA, and surface plasmon resonance (SPR) analyses **(Fig S3C-D, S4, Table S1-3)**. Nbs identified by trypsin and chymotrypsin were comparably high-quality (**Fig S2I**) Overall, 86.2% (CI_95%_: 6.8%), and 90.5% (CI_95%_: 11.5%) true positive Nb binders were verified for GST and HSA respectively (**Fig 2G**). For the challenging PDZ, we used a highly stringent condition for identification; all selected Nb_PDZ_ showed high-affinity binding (**Methods**).

### Accurate large-scale quantification and clustering of Nb proteomes

We evaluated different strategies for accurate classification of Nbs based on affinities. Briefly, antigen-specific HcAbs were affinity isolated from the serum and eluted by the step-wise high-salt gradients, high pH buffers, or low pH buffers (**Methods, Fig S2F-G**). Different HcAbs fractions were accurately quantified by label-free quantitative proteomics (Cox and Mann, 2008; Zhu et al., 2010). The CDR3 peptides (and the corresponding Nbs) were clustered into three groups based on their relative ion intensities (**Fig 3A-B, S3A-B**). Our classification assigns 31% of Nb_GST_ and 47% of Nb_HSA_ into the C3 high-affinity group by the high pH method (**Fig 3C**). We expressed many Nb_GST_ with unique CDR3 sequences from each cluster and measured their affinities by ELISA and SPR (R_2_ = 0.85, **Fig 3D, Table S1**) to evaluate different fractionation methods. Compared with low-pH and high-salt gradient methods, we found that step-wise, high pH-based fractionation methods coupled to quantitative LC/MS can more accurately and reproducibly classify Nbs into three affinity groups. Clusters 1 and 2 (C1, C2) generally having low (μM, C1) and mediocre (dozens of nM, C2) affinities, and C3, 40% of which is represented by sub-nM binders with different binding kinetics (**Fig 3E, 3H, Fig S3C-D**).

**Fig 3.**
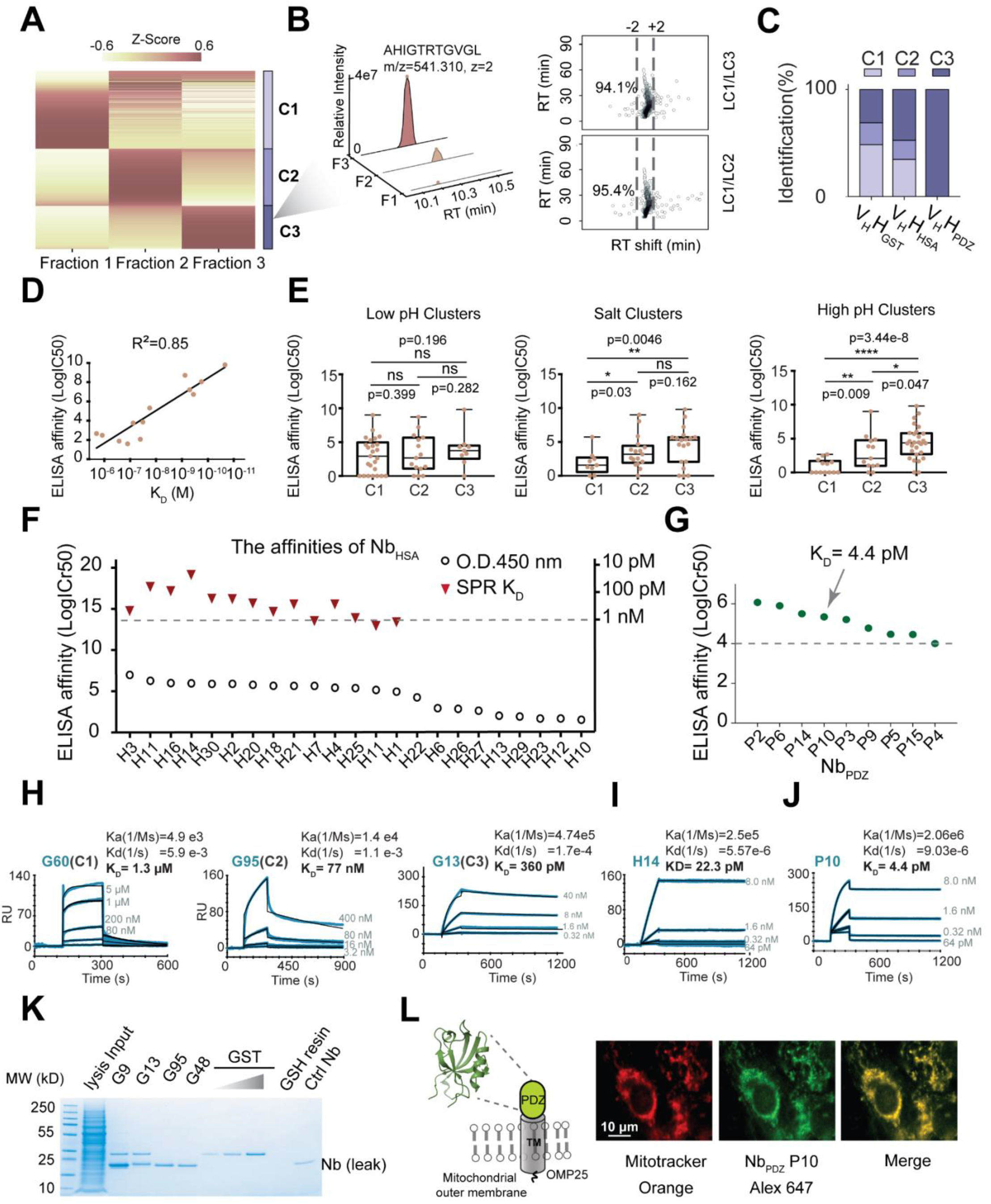
Classification and characterization of antigen-specific Nb repertoires. A) Label-free MS quantification and heat map analysis of CDR3_GST_ fingerprints by chymotrypsin. B) Reproducibility and precision of label-free CDR3_GST_ peptide quantifications by chymotrypsin. C) Percentages of different Nb affinity clusters that were classified by quantitative proteomics. D) Correlation analysis of Nb ELISA affinity (LogIC50 of O.D. 450nm) with SPR K_D_. E) Boxplots of ELISA affinities of different Nb clusters. The p values were calculated based on the student's t test. * indicates a p value of < 0.05, ** indicates < 0.01, *** indicates < 0.001, **** indicates < 0.0001, ns indicates not significant. F) A plot summarizing ELISA affinities of 25 Nb_HSA_ (circles). K_D_ affinities of the top 14 ranked Nbs by ELISA were measured by SPR (red triangles). G) A plot summarizing the ELISA affinities of 11 soluble Nb_PDZ_. H) SPR kinetics analysis of representative Nb_GST_ from three different affinity clusters. I) The SPR kinetics measurement of Nb_PDZ_ P10. J) Immunoprecipitations of GST (1nM concentration) by different Nbs-coupled dynabeads and GSH resin. K) Schematic of the PDZ domain of the mammalian mitochondrial outer membrane protein 25. Fluorescence microscopic analysis of Nb_PDZ_ P10. The Nb was conjugated by Alexa Fluor 647 for native mitochondrial immunostaining of the COS-7 cell line. Mitotracker was used for positive control.

To further verify this result, we purified a random set of 25 Nb_HSA_ (with divergent CDR3s) from C3, and ranked their ELISA affinities (**Fig 3F**, **Table S2**). The top 14 Nb_HSA_ were selected for SPR measurements, in which 11 have dozens to hundreds of pM affinities with different binding kinetics. The remaining 3 Nb _HSA_ demonstrated single-digit nM K_D_’s (**Fig S4A**). We purified 13 soluble Nb_PDZ_ and confirmed their high affinity by ELISA and immunoprecipitation (**Fig 3G**, **S4B**, **Table S3**). We measured the K_D_ of a representative, highly soluble Nb_PDZ_ P10, to be 4.4 pM (**Fig 3I**). We further positively evaluated our high-affinity Nbs for immunoprecipitation (Nb_GST_) and fluorescence imaging (Nb_PDZ_) of native mitochondria (**Fig 3J-K**). Our method enables accurate, large-scale classification of Nb proteomes. Importantly, our results unraveled an unprecedented complexity and diversity of high-affinity circulating Nb repertoires for antigen binding.

### Exploring the mechanisms of Nb affinity maturation

Reliable classifications of the antigen-specific Nbs prompted us to investigate the physicochemical properties that distinguish high-affinity and low-affinity binders towards the same antigen. Shorter CDR3s with distinct distributions were found for high-affinity Nb_HSA_ and Nb_GST_ (**Fig 4A**, **S5A**), likely lowering the entropy for antigen binding (Rini et al., 1992). A significant increase of pIs was also observed with some differences between antigen-specific repertoires: for example, CDR3_HSA_ was primarily responsible for the increase for Nb_HSA_, while CDR1_GST_ and CDR2_GST_ contributed more to the increase of Nb_GST_ (**Fig 4B**, **S5B**). It appears that depending on the epitope properties, Nbs explore different CDR combinations to achieve specific, high-affinity binding.

**Fig 4.**
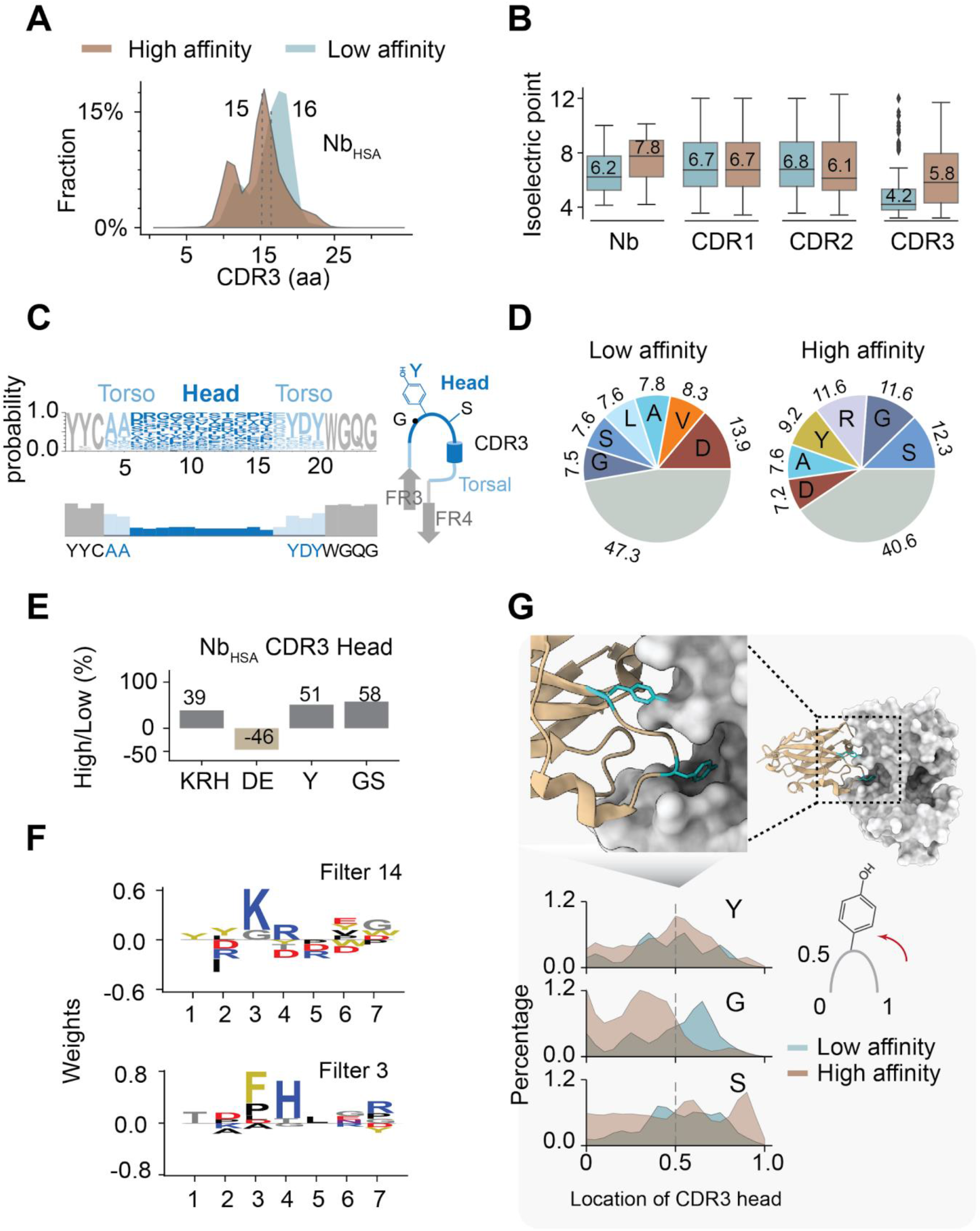
Mechanisms of Nb affinity maturation. A) Distributions of CDR3 lengths of high-affinity (brown) and low-affinity (steel) Nb_HSA_. B) Comparisons of the pI of low- and high- affinity Nbs and different CDRs. C) Alignment of CDR3 sequences based on a random selection of 1,000 unique CDR3 sequences with the identical length of 15 residues.D) Pie charts of the amino acid compositions of the CDR3 heads of low- and high- affinity Nb_HSA_. The 6 most abundant residues are shown. E) The relative changes of abundant amino acids on CDR3 heads of Nb_HSA_. F) Sequence logo of two representative convolutional CDR3 filters (Filter 14 for high-affinity Nb_HSA_; filter 3 for low-affinity Nb_HSA_) learned by a deep learning model. G) Comparisons of the relative abundance of Y, G and S on the CDR3 heads between high-affinity and low-affinity Nb_HSA_. Their relative abundances are plotted as a function of the relative position of the respective residues. A representative structure (PDB: 5F1O) showing two tyrosines on the Nb CDR3 head are inserted into the deep pockets of the antigen.

A CDR3 fingerprint sequence can be divided into a central “head” region containing the highest sequence variability, and a lower specificity adjacent “torso” region (Finn et al., 2016; Lam et al., 2009) (**Fig 4C**). Specific residues, including aspartic acid and arginine (forming strong electrostatic interactions), aromatic tyrosine, small and flexible glycine, and serine, and hydrophobic alanine and leucine were enriched on the heads (**Fig 4D-E**, **S3C-E**). We identified two main features for high-affinity binders: 1) significant alterations of charged residues, likely crucial for optimizing the electrostatic complementarity between the paratopes and corresponding epitopes, and 2) an increase in the number of tyrosines on the CDR3 heads. Interestingly, for high-affinity Nb_HSA_, tyrosine is more frequently found at the center of the CDR3 loops, presumably inserting its bulky, aromatic side-chain into the specific pocket(s) on the HSA epitopes (Li et al., 2016; Peng et al., 2014). Glycine and serine tend to be placed away from the CDR3 center, providing additional flexibilities and facilitating the orientation of the tyrosine side chain in the pocket (**Fig 4G**). These results were supported by the correlation analysis between the number of these residue groups and ELISA affinities. For example, we found a significant negative correlation between the number of negatively charged residues of Nb_HSA_ and ELISA affinity (Pearson r = −0.48) and a positive association for tyrosine and glycine (Pearson r = 0.52, **Fig S5F**). Correlation plots of ELISA affinities and the number of specific residues revealed different slopes (−1.0 for negative charges and 0.6 for tyrosine and glycine), potentially reflecting the differences in contributions to Nb affinity between electrostatic and hydrophobic interactions. For Nb_GST_, we found a positive correlation between CDR2 charges and the affinity (**Fig S5G**).

We developed a deep learning model and trained the network using the sequences of Nbs with their corresponding affinity clusters as labels for a complimentary analysis. We extracted features unique to each group of Nbs once the network learned the sequence patterns (filters) that enable affinity classification (92% accuracy, **Methods**). Deep learning revealed a pattern of consecutive lysine and arginine, tyrosine, and glycines as the most informative CDR3 feature for high-affinity Nb_HSA_ (**Methods**, **Fig 4F**, **S5H**). For low-affinity binders, the most informative filter has a preference for phenylalanine, histidine, and two consecutive aspartic acids. A quick comparison of the consensus sequences reveals a tendency for successive pairs of positive charges in the heads of high-affinity binders and consecutive pairs of negative charges in the heads of low-affinity binders.

### The landscapes of antigen-engaged Nb proteomes revealed by integrative structure methods

High-throughput docking and clustering analysis of 34,972 unique Nb_HSA_ revealed three dominant, non-linear/conformational epitopes (E1, E2, and E3) on HSA (**Fig 5A**). We cross-linked 19 HSA-Nb complexes using complementary cross-linkers of amine-specific disuccinimidyl suberate (DSS), and a heterobifunctional carbodiimide crosslinker (EDC) that reacts with both lysine and acidic residues (aspartic acid and glutamic acid) (Kim et al., 2018; Shi et al., 2014). Cross-linking mass spectrometry confirmed the docking results (5%, 60%, 20%, E1, E2, and E3, respectively) and identified two additional minor epitopes (**Fig 5B-D**). 92% of cross-links were satisfied by the models with a median RMSD of 5.6 Å (**Fig 5E-F**). To further verify the cross-link models, we introduced a single point mutation E400R on HSA with minimal impact on the overall structure as predicted by mCSM (change of stability ΔΔG=-0.1) (Pires et al., 2014). E400R potentially disrupts a salt bridge formed between the most abundant E2 epitope and arginine on the Nb CDR3 (**Fig 5G**). The mutation significantly weakened 26% of the interactions that we tested using ELISA, confirming that E2 is a *bona fide* dominant epitope.

**Fig 5.**
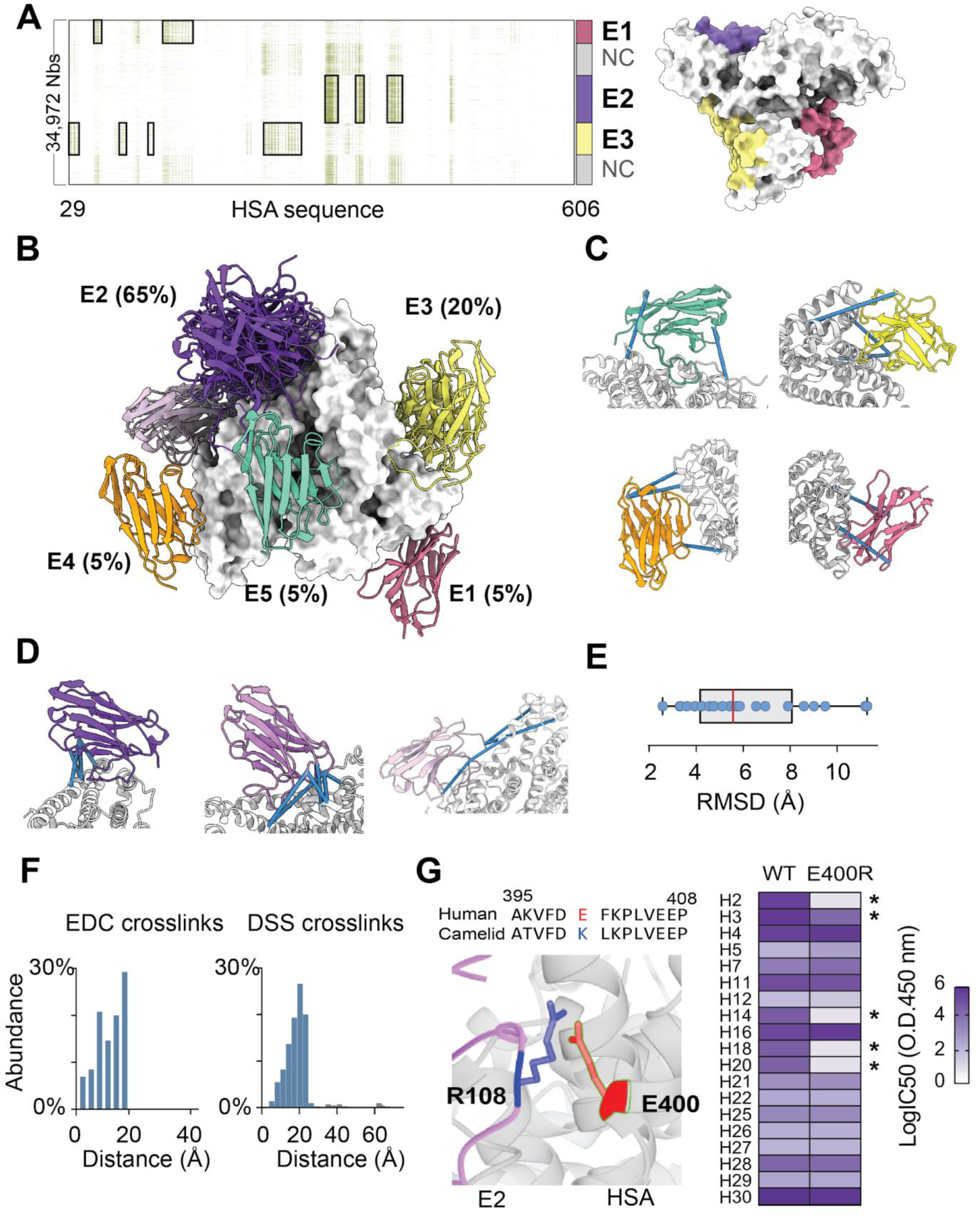
The structural landscapes of HSA-specific Nb proteomes. A) The heatmap of the major epitopes mapped by structural docking and the surface representation of the three dominant HSA epitopes. HSA are presented in gray. E1, E2 and E3 are in pale red, purple and yellow, respectively. B) The HSA epitopes and their fractions (%) based on converged cross-link models (E1: residues 57-62, 135-169; E2: 322-331, 335, 356-365, 395-410; E3: 29-37, 86-91, 117-123, 252-290; E4: 566-585, 595, 598-606 and E5:188-208, 300-306, 463-468). C) Representative cross-link models of HSA-Nb complexes. The best scoring models were presented. Satisfied DSS or EDC cross-links are shown as blue sticks. D) Representative cross-link models of HSA-Nb complexes on the dominant epitope E2, which contains three sub-epitopes (variants that share certain E2 residues but do not completely overlap) that were indicated in different purple colors. E) A plot of the RMSDs (room-median-square-deviations) of HSA-Nb cross-link models. F) Bar plots showing the % of the GST-Nb cross-link satisfactions (EDC crosslinks <25 angstrom and DSS crosslinks < 30 angstrom). G) Left: A putative salt bridge between glutamic acid 400 (HSA) and arginine 108 of a Nb CDR3 is presented. The local sequence alignment between HSA and camelid albumin is shown. Right: ELISA affinity screening (heatmap) of 19 different Nbs for binding to wild type HSA and the point mutant (E400R). * indicates decreased affinity.

Using this approach, we identified and experimentally verified three dominant epitopes for Nb_GST_ (**Fig 6A**): E1 (18.75%) and E3 (50%) contain charged surface patches. E2 (31.25%) overlapped with the GST dimerization cavity (**Fig 6B, 6E**), where the CDR3 loops of Nbs were inserted for binding. 91% of cross-links were satisfied by the models with a median RMSD of 6 Å (**Fig 6C-D**). Two epitopes were identified for PDZ (**Fig S6A**). E2_PDZ_ had a structured surface consisting of an α helix and two β-strands and was hypothesized to be the dominant epitope. We then produced a double mutant PDZ (R46E: K48D) that, according to our models, disrupts two important salt bridges for high-affinity interaction (**Fig S6B**), and evaluated the mutant binding by ELISA. The majority (8/11) of the Nb_PDZ_ that we selected exhibited significantly decreased or no affinity for the mutant, confirming that E2 is the dominant epitope (**Fig S6C**).

**Fig 6.**
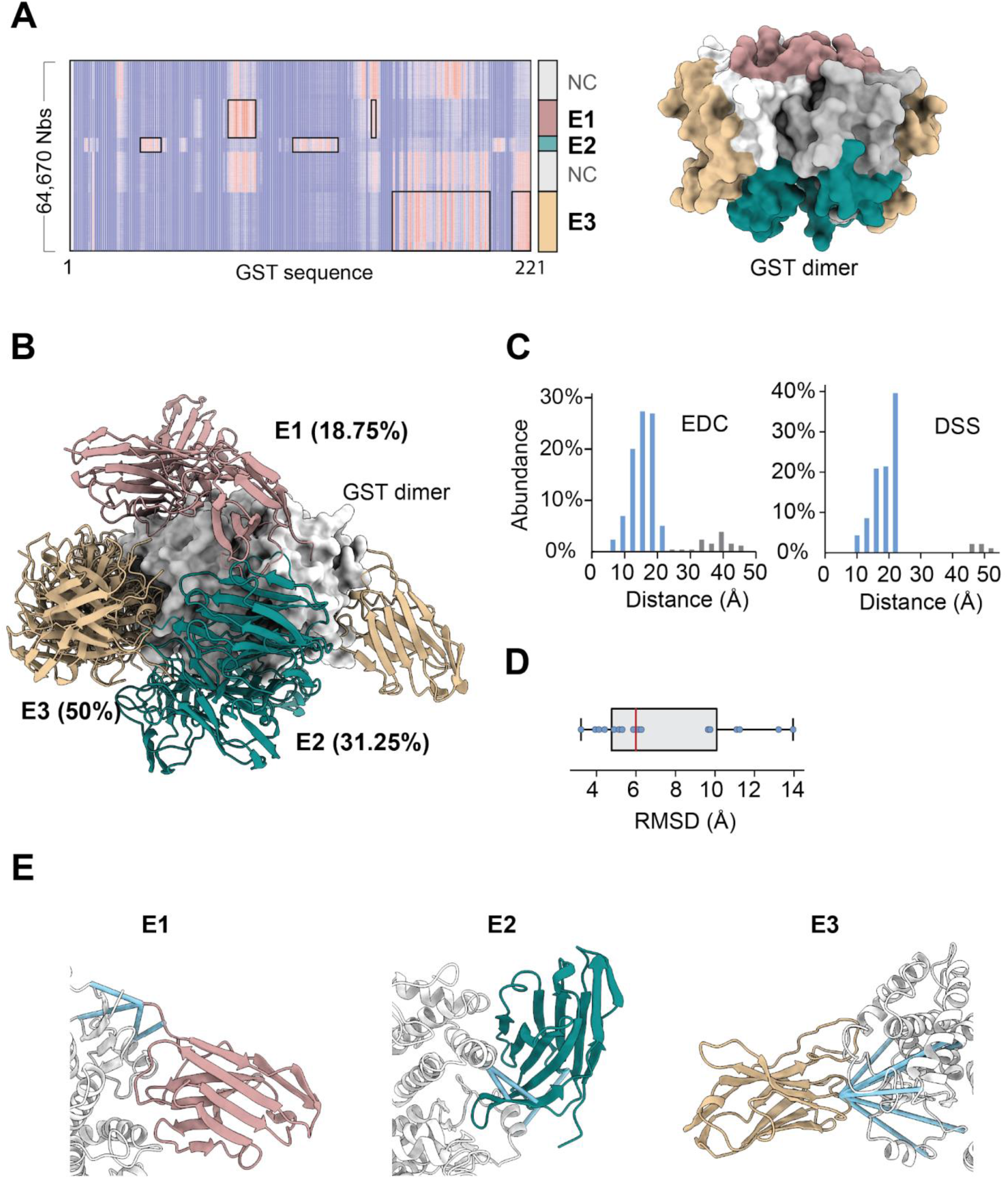
Hybrid structural analysis of GST-Nb complexes. A) Heatmap analysis of structural docking of 64,670 GST-Nb complexes showing three converged epitopes (E1: 75-88, 143-148; E2: 33-43, 107-127; E3: 158-200, 213-220). Surface representations of the GST dimer were shown. GST dimers were presented in gray. E1, E2 and E3 were in rosy brown, teal and tan respectively. B) GST epitopes and their abundances (%) based on converged cross-link models were shown with corresponding colors. C) Bar plots showing the % of the GST-Nb cross-link satisfactions. D) A plot of the RMSD of GST-Nb cross-link models. E) Representative cross-link models of different GST-Nb complexes. Blue sticks indicate DSS or EDC cross-links that are within the respective distance thresholds (i.e., 25 and 20 angstrom, respectively).

### The mechanisms that underlie antigen binding by high-affinity Nbs

We have developed a high-throughput platform for the analysis of circulating antigen-engaged Nbs. Large repertoires of diverse and high-affinity Nb antibodies were discovered, quantified, affinity-classified, and structurally characterized. Our analysis revealed a remarkable specificity, robustness, versatility, diversity of the antigen-engaged circulating Nb repertoires (**Fig 7A**). The specificity was well exemplified in HSA binding, where Nbs evolved to specifically recognize HSA surface pockets that differ in physicochemical properties (i.e., pI and hydropathy) between “self” endogenous llama albumin and “non-self” HSA (sharing 87% sequence similarity) to avoid an autoimmune response (**Fig 7B**).

**Fig 7:**
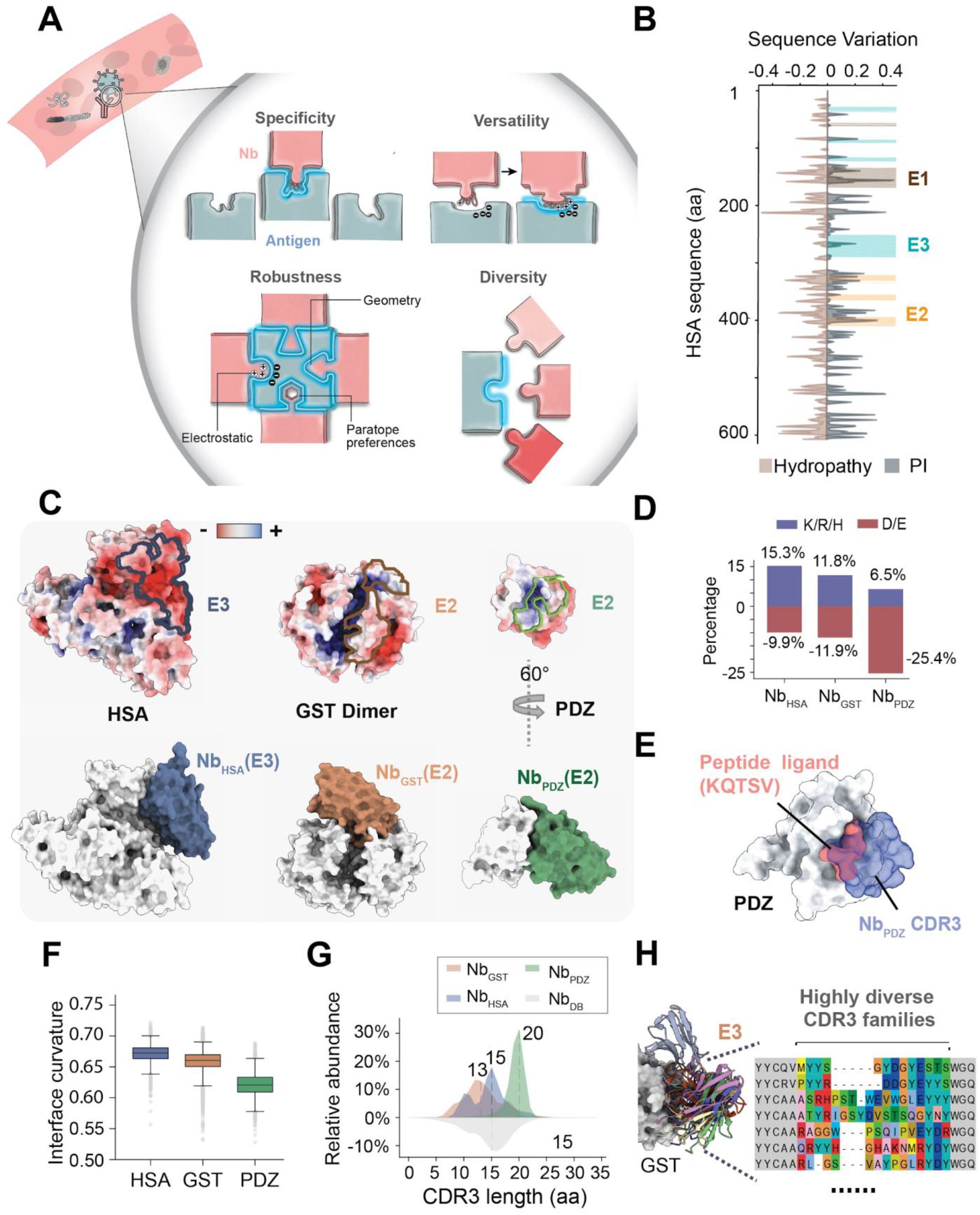
Mechanisms that underlie antigen binding by a large repertoire of high-affinity Nbs. A) A schematic cartoon model for antigen binding by Nbs. B) The sequence variations of pI and hydropathy between human and camelid serum albumins. C) Surface representations showing co-localization of electrostatic potential surfaces with epitopes on three antigens. D) Bar plots showing percentages of charged residues on CDR3 heads of different Nbs. E) Surface representations showing the shared binding region between natural peptide ligand (KQTSV, PDB:1BE9) and a Nb_PDZ_ CDR3 on PDZ. The peptide ligand was colored in salmon and CDR3 was transparent and colored in slate. F) Boxplots showing the relative interface curvature of different antigens based on high-throughput structural docking of antigen-Nb complexes (lower value corresponds to flatter interfaces). G) Plot comparisons of the CDR3 lengths of different high-affinity Nbs. The curve was smoothed with a gaussian function. H) Cartoon representations of the cross-link models of a diverse set of high-affinity Nb_GST_ that bind to the same epitope (E3) on GST. The CDR3 sequence alignment was shown.

The robustness of antigen-Nb interactions is achieved by optimizing electrostatic and shape complementarity (Lawrence and Colman, 1993; McCoy et al., 1997). We found that the most abundant epitopes are generally enriched with charged residues (**Fig 7C, upper panel**), while complementary charges were identified on the CDR3s of the corresponding Nbs (**Fig 7D, S7A, S7D**). Consistently, high-affinity binders have higher numbers of charged residues compared to low-affinity ones. While we observed preference of Nbs to concave-shaped epitopes, they can also bind flat surfaces (**Fig 7C, lower panel**). Moreover, the shape and charge complementarity are increased during affinity maturation through enrichment and redistribution of specific groups of “hot-spot” paratope residues (charged residues, tyrosine, and small hydrophobic residues).

The versatility of the immune system was revealed by analyzing the response to the weakly immunogenic PDZ domain. Here, specific Nbs evolved unusually long CDR3 loops (20 residues by a median, **Fig 7G**) enriched with aspartic acid, glycine, and serine, which account for half of the CDR3 residues (**Fig S7A, S7D**). The long CDR3 loops wrap around the relatively shallow (**Fig 7F**) yet positively charged epitope to create extensive electrostatic and hydrophobic interactions. Interestingly, this epitope overlaps with a conserved ligand-binding motif shared among numerous PDZ interacting proteins (**Fig 7E**) (Doyle et al., 1996; Sheng and Sala, 2001). Nb_PDZ_ are acidic with a median pI of 4.9 pushing the envelope of their physicochemical properties to achieve >100,000-fold higher affinity (**Fig 3I, S7B**) than the natural intracellular ligands (typically in μM affinity) (Niethammer et al., 1998). Despite their acidic nature, Nb_PDZ_ did not seem to appreciably alter hydropathy, due to the compensation of hydrophobic residues(**Fig S7C**). Here we took a snapshot of this fascinating rapid evolution of protein-protein interaction, revealing a convergent force that drives this process.

## Discussion

The unprecedented diversity of high-affinity Nbs for antigen engagement was among the most significant observations. Thousands of exceptionally diverse, specific CDR3 Nb families were identified for a variety of dominant epitopes on the antigens. Even within an epitope, there exists a large cohort of different binders (**Fig 7H**). It has been shown that immunodominance may improve the efficiency of the immune response (Akram and Inman, 2012). Here the coexistence of numerous high-affinity epitope antibodies in the serum, seemingly redundant, may facilitate robust, synergistic binding and clearance of challenging pathogens such as viruses that constantly mutate to escape from the immune surveillance (Sanjuan et al., 2010; Scheid et al., 2011; Wei et al., 2003). Our big data thus sheds light on the epic landscapes of mammalian humoral immunity and uncovers a powerful potential mechanism by which the immune response has evolved to enact on this never-ending race.

We envision that our technologies will find broad utility in challenging biomedical applications such as cancer biology, brain research, and virology. The high-quality Nb dataset and the structural models presented here may serve as a blueprint to study antibody-antigen and protein-protein interactions that orchestrate biological processes.

## Supplemental Tables 1-3

Table S1 Summary of GST-Nbs

Table S2 Summary of HSA-Nbs

Table S3 Summary of PDZ-Nbs

**Supplemental Item 1**(cross-link model files)

The proteomics data of chemical crosslink and mass spectrometric analysis (CX-MS) analysis has been deposited into MassIVE data repository.

## STAR* Methods

### KEY RESOURCES TABLE

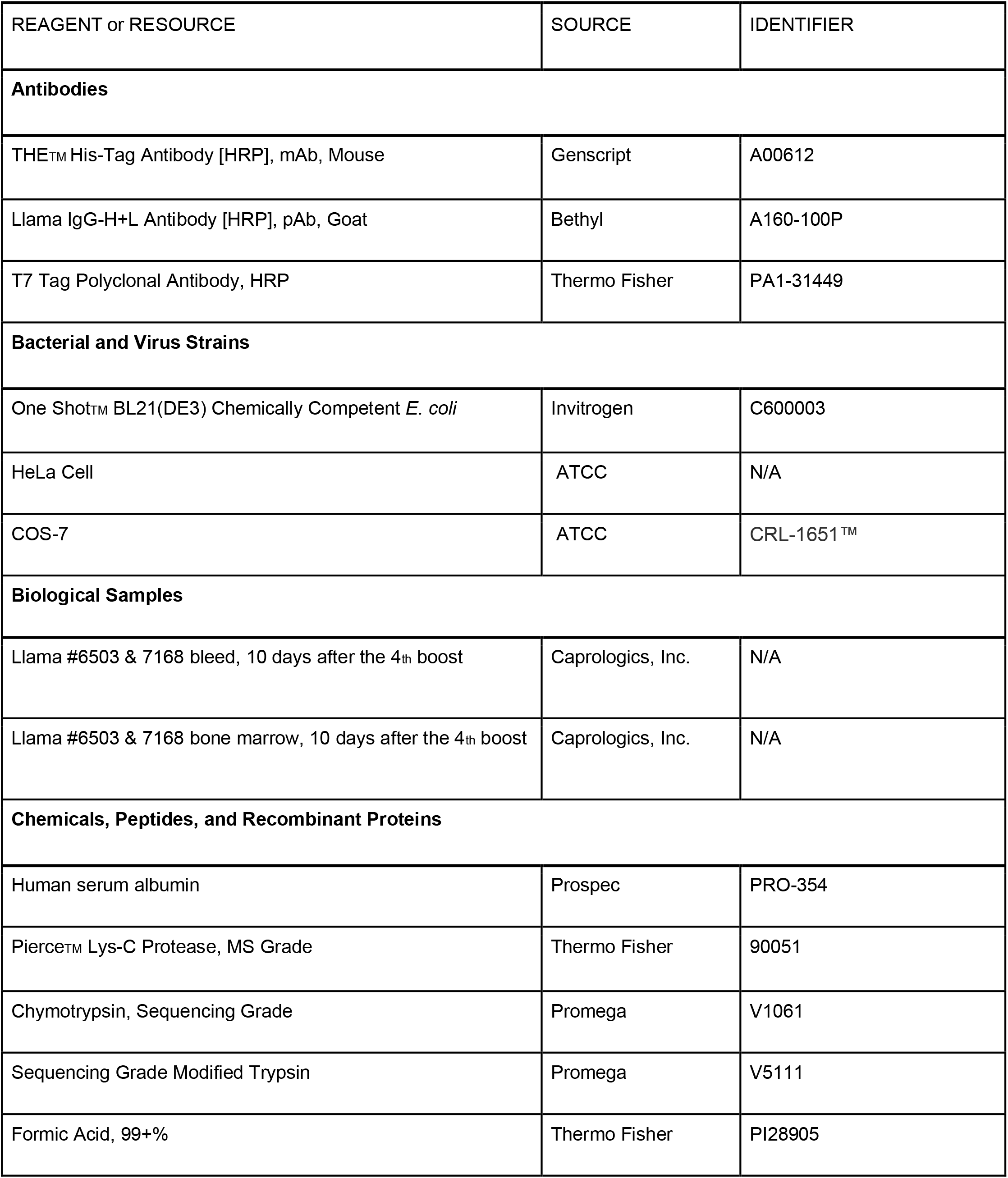

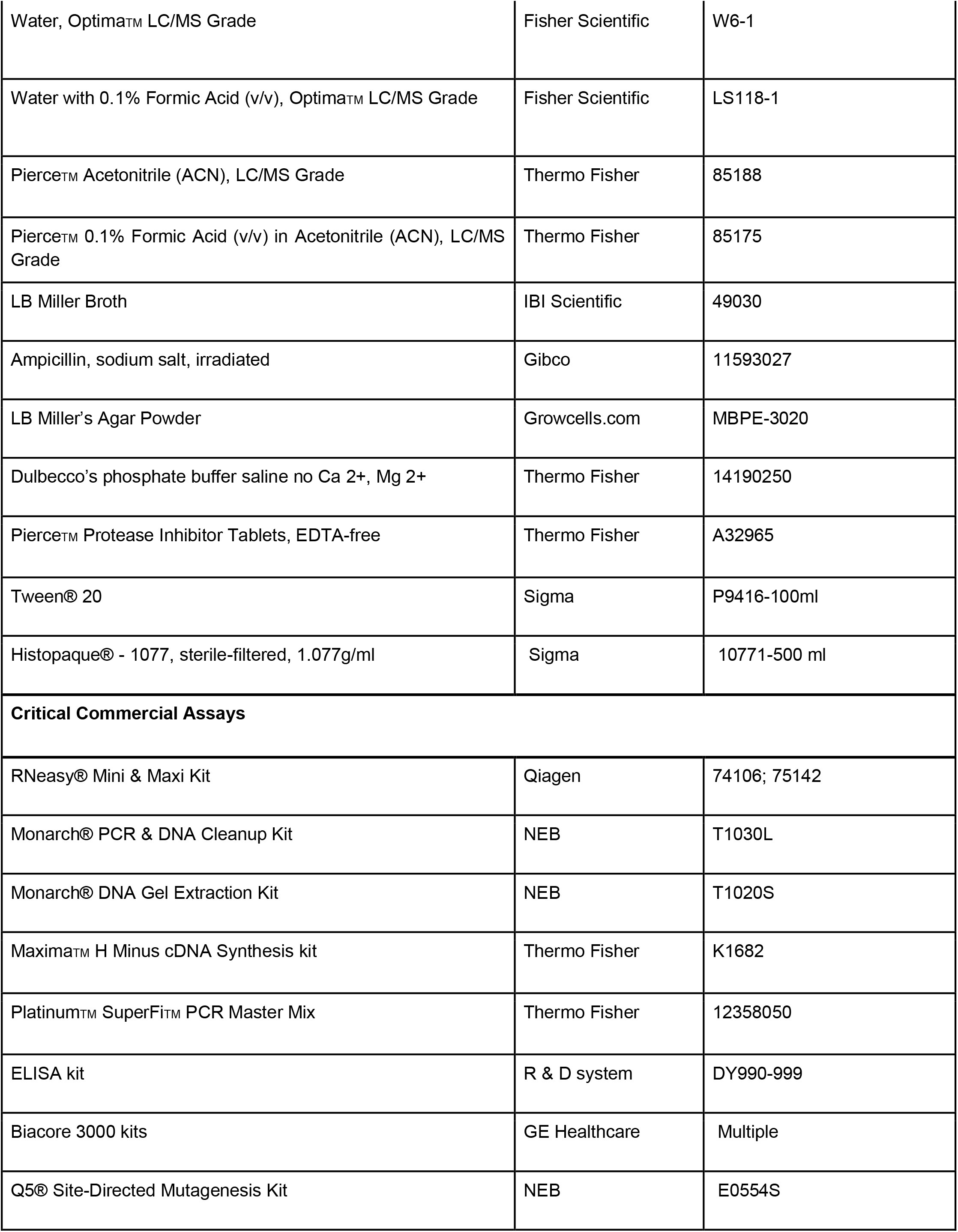

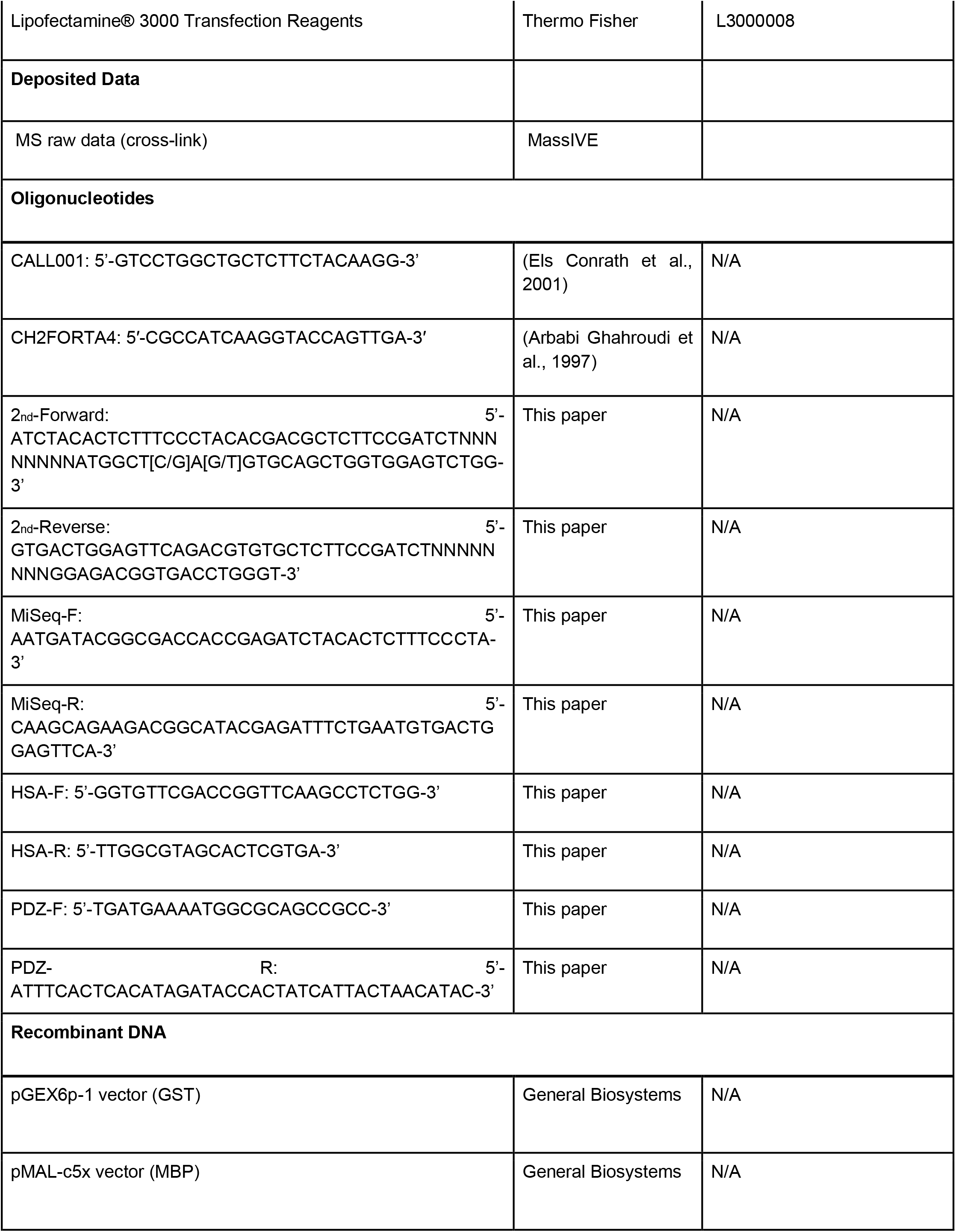

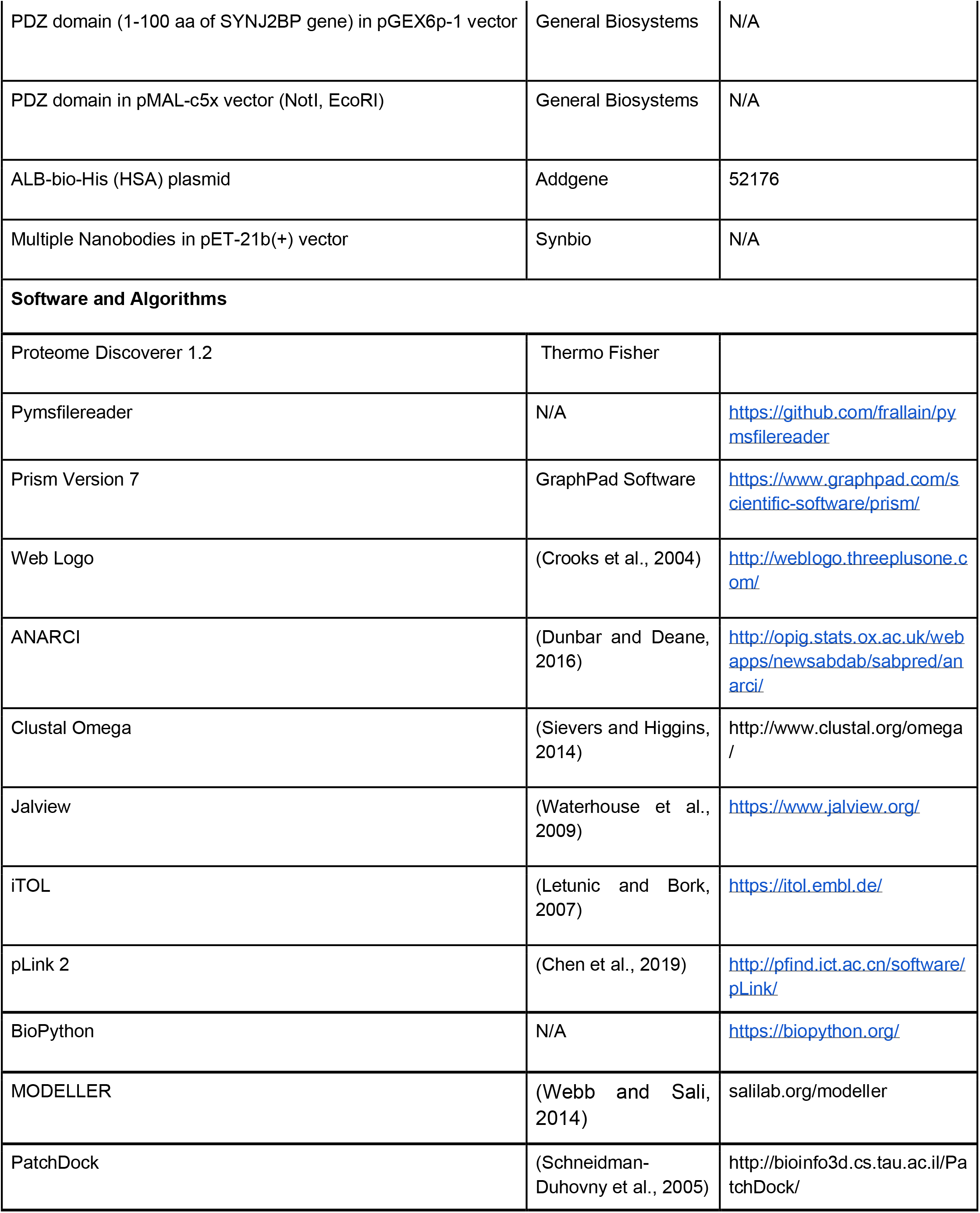

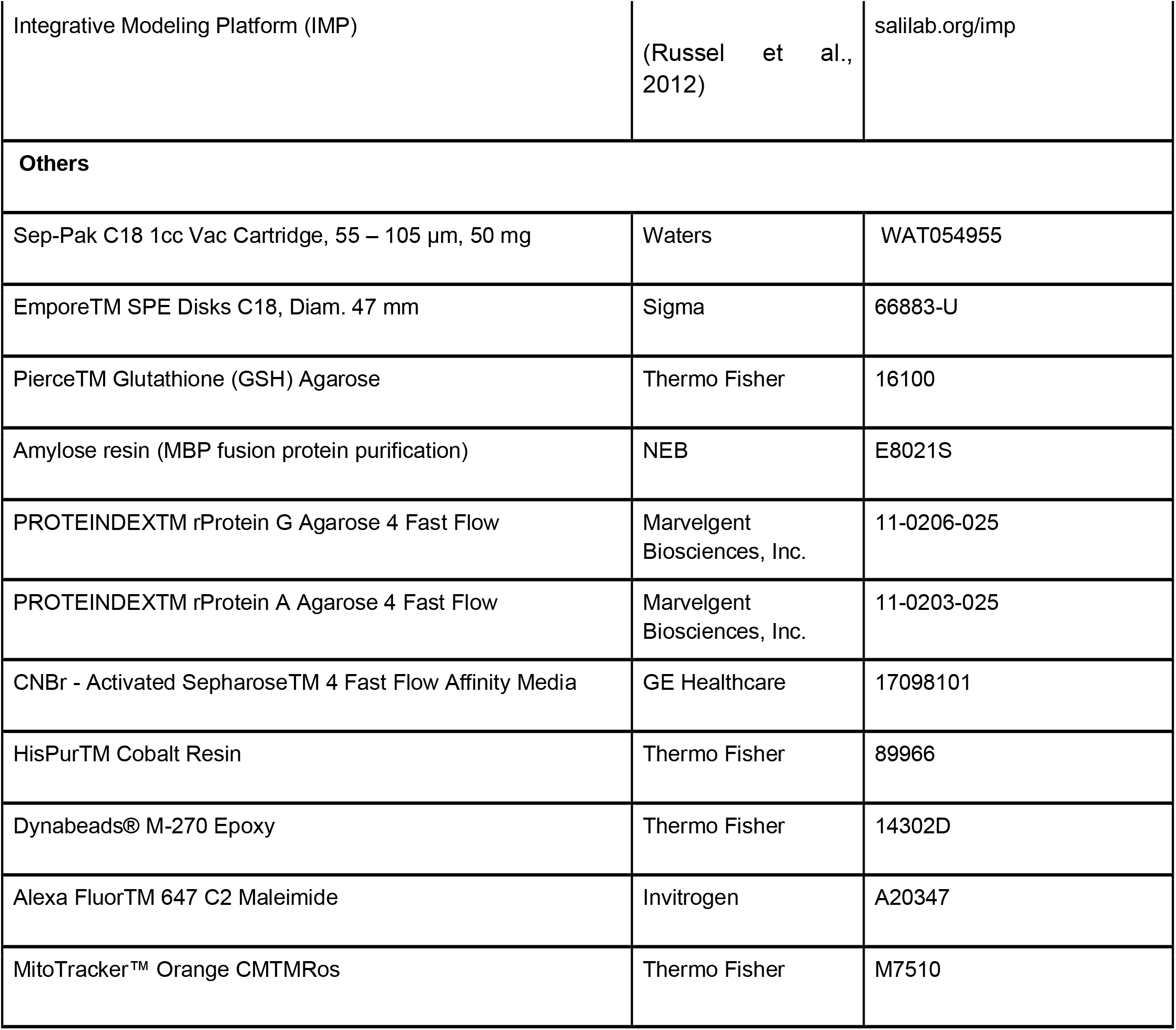

### Animal immunization

Two Llamas were respectively immunized with HSA, and a combination of GST and GST fusion PDZ domain of Mitochondrial outer membrane protein 25 (OMP25) at the primary dose of 1 mg, followed by three consecutive boosts of 0.5 mg every 3 weeks. The bleed and bone marrow aspirates were extracted from the animals 10 days after the last immuno-boost. All the above procedures were performed by Capralogics, Inc. following the IACUC protocol.

### mRNA isolation and cDNA preparation

~ 1 - 3 ×10^9^ peripheral mononuclear cells were isolated from 350 ml immunized blood and 5 - 9 ×10^7^ plasma cells were isolated from 30 ml bone marrow aspirates using Ficoll gradient (Sigma). The mRNA was isolated from the respective cells using RNeasy kit (Qiagen) and was reverse-transcribed into cDNA using Maxima™ H Minus cDNA Synthesis Master Mix (Thermo). Camelid IgG heavy chain cDNA sequences from the variable domain to the CH2 domain were specifically amplified using primers CALL001 and CH2FORTA4 (Arbabi Ghahroudi et al., 1997). The VHH genes that lack CH1 domain were separated from conventional IgG by DNA gel electrophoresis and were subsequently re-amplified from framework 1 to framework 4 using the 2nd-Forward and 2nd-Reverse. The random 8-mers replacing adaptor sequences were added to aid in cluster identification for Illumina MiSeq. The amplicon of the second PCR (approximately 450-500 bp) was purified using Monarch PCR clean up kit. The final round of PCR with primer MiSeq-F and MiSeq-R was performed to add P5 and P7 adapters with the index before MiSeq sequencing.

### Next generation sequencing by Illumina Miseq

Sequencing was performed based on the Illumina MiSeq platform with the 300 bp paired-end model. More than 32.4 million reads were generated with a median phred score 34 corresponding to ~ 0.1% error rate. Read QC tool in FastQC v0.11.8 (http://www.bioinformatics.babraham.ac.uk/projects/fastqc/) was used for quality check and control of the FASTQ data. Raw Illumina reads were processed by the software tools from the BBMap project (https://github.com/BioInfoTools/BBMap/). Duplicated reads and DNA barcode sequences were removed successively before converting the nucleotide sequences into amino acid sequences. Eventually 13.6 million high-quality unique protein sequences were generated. The complementary determining regions were annotated according to AbM CDR definition (Abhinandan and Martin, 2008).

### Isolation and biochemical fractionation of VHH antibodies from immunized sera

Approximately 175 ml of plasma was isolated from 350 ml of immunized blood by Ficoll gradient (Sigma). Camelid single-chain V_H_H antibodies were isolated from the plasma supernatant by a two-step purification procedure using protein G and protein A sepharose beads (Marvelgent), acid-eluted, before neutralized and diluted in 1xPBS buffer to a final concentration of 0.1- 0.3 mg/ml. To purify antigen-specific V_H_H antibodies, the GST or HSA-conjugated CNBr resin was incubated with the VHH mixture for 1 hr at 4°C and extensively washed with high salt buffer (1xPBS and 350 mM NaCl) to remove non-specific binders. Specific V_H_H antibodies was then released from the resin by using one of the following elution conditions: alkaline (1-100 mM NaOH, pH 11, 12 and 13), acidic (0.1 M glycine, pH 3, 2 and 1) or salt elution (1M – 4.5 M MgCl2 in neutral pH buffer). For purification of PDZ-specific V_H_H, a fusion protein of MBP-PDZ (where the maltose binding protein/MBP was fused to the N terminus of PDZ domain to avoid steric hindrance of the small PDZ after coupling) was produced and was used as the affinity handle. MBP coupled resin was used for control (See supplementary Fig 2). All the eluted V_H_Hs were neutralized and dialyzed into 1x DPBS separately prior to proteomics analysis.

### Proteolysis of Antigen Specific Nbs and Nanoflow Liquid Chromatography coupled to Mass spectrometry (nLC/MS) Analysis

For GST and HSA V_H_Hs, each elution was processed separately according to the following protocol. For PDZ specific VHHs, only the most stringent biochemical elutes (i.e., pH 13, pH 1, MgCl_2_ 3M and 4.5M) and the respective nonspecific MBP binders (negative controls) from different fractions were pooled for proteolysis. For instance, For PDZ-specificV_H_Hs that were eluted by pH13 buffer, we pooled non-specific MBP binding Nbs from pH 11, pH12 and pH13 fractions for negative control to improve the stringency of our downstream LC/MS quantification. VHHs were reduced in 8M urea buffer (with 50 mM Ammonium bicarbonate, 5 mM TCEP and DTT) at 57°C for 1hr, and alkylated in the dark with 30 mM Iodoacetamide for 30 mins at room temperature. The alkylated sample was then split into two and in-solution digested using either trypsin or chymotrypsin. For trypsin digestion samples, 1:100 (w/w) trypsin and Lys-C were added and digested at 37°C overnight, with additional 1:100 trypsin the other morning for 4 hrs at 37°C water bath. For chymotrypsin digestion samples, 1:50 (w/w) chymotrypsin was added and digested at 37 °C for 4 hrs. After proteolysis, the peptide mixtures were desalted by self-packed stage-tips or Sep-pak C18 columns (Waters) and analyzed with a nano-LC 1200 that is coupled online with a Q Exactive™ HF-X Hybrid Quadrupole Orbitrap™ mass spectrometer (Thermo Fisher). Briefly, desalted Nb peptides were loaded onto an analytical column (C18, 1.6 μm particle size, 100 Å pore size, 75 μm × 25 cm; IonOpticks) and eluted using a 90-min liquid chromatography gradient (5% B–7% B, 0–10 min; 7% B–30% B, 10–69 min; 30% B–100% B, 69 – 77 min; 100% B, 77 - 82 min; 100% B - 5% B, 82 min - 82 min 10 sec; 5% B, 82 min 10 sec - 90 min; mobile phase A consisted of 0.1% formic acid (FA), and mobile phase B consisted of 0.1% FA in 80% acetonitrile (ACN)). The flow rate was 300 nl/min. The QE HF-X instrument was operated in the data-dependent mode, where the top 12 most abundant ions (mass range 350 – 2,000, charge state 2 - 8) were fragmented by high-energy collisional dissociation (HCD). The target resolution was 120,000 for MS and 7,500 for tandem MS (MS/MS) analyses. The quadrupole isolation window was 1.6 Th and the maximum injection time for MS/MS was set at 80 ms.

### Nb DNA synthesis and cloning

Nb genes were codon-optimized for expression in *Escherichia coli* and the nucleotides were *in vitro* synthesized (Synbiotech). After verification by Sanger sequencing, the Nb genes were cloned into a pET-21b (+) vector at BamHI and XhoI (for Nb_GST_), or EcoRI and NotI restriction sites (Nb_PDZ_ and Nb_HSA_).

### Purification of recombinant Proteins

DNA constructs were transformed into BL21(DE3) competent cells according to manufacturer's instructions and plated on Agar with 50 μg/ml ampicillin at 37 °C overnight. A single colony was inoculated in LB medium with ampicillin for overnight culture at 37 °C. The culture was then inoculated at 1:100 (v/v) in fresh LB medium and shaked at 37 °C until the O.D.600 nm reached 0.4-0.6. GST, GST-PDZ and Nbs were induced with 0.5 mM of IPTG while MBP and MBP-PDZ were induced with 0.1 mM of IPTG. The inductions were performed at 16°C overnight. Cells were then harvested, briefly sonicated and lysed on ice with a lysis buffer (1xPBS, 150 mM NaCl, 0.2% TX-100 with protease inhibitor). After lysis, soluble protein extract was collected at 15,000 x g for 10 mins. GST and GST-PDZ were purified using GSH resin and eluted by glutathione. MBP (maltose binding protein) and MBP-PDZ fusion protein were purified by using Amylose resin and were eluted by maltose according to the manufacturer's instructions. Nbs were purified by His-Cobalt resin and were eluted using imidazole. The eluted proteins were subsequently dialyzed in the dialysis buffer (e.g., 1x DPBS, pH 7.4) and stored at −80 °C before use.

### Nb immunoprecipitation assay

After Nb induction and cell lysis, the cell lysates were run on SDS-PAGE to estimate Nb expression levels. Recombinant Nbs in the cell lysis were diluted in 1x DPBS (pH 7.4) to a final concentration of ~ 5 μM (for Nb_GST_) and ~ 50 nM (for Nb_PDZ_). To test the specific interactions of Nbs with antigens, different antigens were coupled to the CNBr resin. Inactivated or MBP-conjugated CNBr resin was used for control. Antigen coupled resins or control resins were incubated with Nb lysates at 4°C for 30 mins. The resins were then washed three times with a washing buffer (1x DPBS with 150 mM NaCl and 0.05% Tween 20) to remove nonspecific bindings. Specific antigen bound Nbs were then eluted from the resins by a hot LDS buffer containing 20 mM DTT and ran on SDS-PAGE. The intensities of Nbs on the gel were compared between antigen specific signals and control signals to derive the false positive binding.

### ELISA (enzyme-linked immunosorbent assay)

Indirect ELISA was carried out to evaluate the camelid immune response of an antigen and to quantify the relative affinities of antigen-specific Nbs. An antigen was coated onto a 96-well ELISA plate (R&D system) at an amount of approximately 1-10 ng per well in a coating buffer (15 mM sodium carbonate, 35 mM sodium bicarbonate, pH 9.6) overnight at 4°C. The well surface was then blocked with a blocking buffer (DPBS, 0.05% Tween 20, 5% milk) at room temperature for 2 hours. To test an immune response, the immunized serum was serially 5-fold diluted in the blocking buffer. The diluted sera were incubated with the antigen coated wells at room temperature for 2 hours. HRP-conjugated secondary antibodies against llama Fc were diluted 1:10,000 in the blocking buffer and incubated with each well for 1 hour at room temperature. For Nb affinity tests, scramble Nbs that do not bind the antigen of interest were used for negative controls. Nbs of both specific binders for test and scramble negative controls were serially 10-fold diluted from 10 μM to 1 pM in the blocking buffer. HRP-conjugated secondary antibodies against His-tag or T7-tag were diluted 1:5,000 or 1:10,000 in the blocking buffer and incubated for 1 hour at room temperature. Three washes with 1x PBST (DPBS, 0.05% Tween 20) were carried out to remove nonspecific absorbances between each incubation. After the final wash, the samples were further incubated under dark with freshly prepared w3,3',5,5'-Tetramethylbenzidine (TMB) substrate for 10 mins at room temperature to develop the signals. After the STOP solution (R&D system), the plates were read at multiple wavelengths (450 nm and 550 nm) on a plate reader (Multiskan GO,Thermo Fisher). A false positive Nb binder was defined if any of the following two criteria was met: i) the ELISA signal could only be detected at a concentration of 10 μM and was under detected at 1 μM concentration. ii) at 1 μM concentration, a pronounced signal decrease (by more than 10 fold) was detected compared to the signal at 10 μM, while there were no signals could be detected at lower concentrations. The raw data was processed by Prism 7 (GraphPad) to fit into a 4PL curve and to calculate logIC50.

### Nb affinity measurement by SPR

Surface plasmon resonance (SPR, Biacore 3000 system, GE Healthcare) was used to measure Nb affinities. Antigen proteins immobilized on the activated CM5 sensor-chip by the following steps. Protein analytes were diluted to 10-30 μg/ml in 10 mM sodium acetate, pH 4.5, and were injected into the SPR system at 5 μl/min for 420 s. The surface of the sensor was then blocked by 1 M ethanolamine-HCl (pH 8.5). For each Nb analyte, a series of dilution (spanning three orders of magnitude) was injected in HBS-EP+ running buffer (GE-Healthcare) containing 2 mM DTT, at a flow rate of 20- 30 μl/min for 120-180 s, followed by a dissociation time of 5 - 20 mins based on dissociation rate. Between each injection, the sensor chip surface was regenerated with the low pH buffer containing 10 mM glycine-HCl (pH 1.5 - 2.5), or high pH buffer of 20 - 40 mM NaOH (pH 12 - 13). The regeneration was performed with a flow rate of 40-50 μl/min for 30 s. The measurements were duplicated and only highly reproducible data was used for analysis. Binding sensorgrams for each Nb were processed and analyzed using BIAevaluation by fitting with 1:1 Langmuir model or 1:1 Langmuir model with mass transfer.

### Cross-linking and mass spectrometric analysis of antigen-nanobody complex

Different Nbs were incubated with the antigen of interest with equal molarity in an amine-free buffer (such as 1x DPBS with 2 mM DTT) at 4°C for 1 - 2 hours before cross-linking. The amine-specific disuccinimidyl suberate (DSS) or heterobifunctional linker 1-ethyl-3-(3-dimethylaminopropyl) carbodiimide hydrochloride (EDC) was added to the antigen-Nb complex at 1 mM or 2 mM final concentration, respectively. For DSS cross-linking, the reaction was performed at 23°C for 25 mins with constant agitation. For EDC cross-linking, the reaction was performed at 23°C for 60 mins. The reactions were quenched by 50 mM Tris-HCl (pH 8.0) for 10 mins at room temperature. After protein reduction and alkylation, the cross-linked samples were separated by a 4–12% SDS-PAGE gel (NuPAGE, Thermo Fisher). The regions corresponding to the cross-linked species were cut and in-gel digested with trypsin and Lys-C as previously described (Shi et al., 2014; Shi et al., 2015). After proteolysis, the peptide mixtures were desalted and analyzed with a nano-LC 1200 (Thermo Fisher) coupled to a Q Exactive™ HF-X Hybrid Quadrupole-Orbitrap™ mass spectrometer (Thermo Fisher). The cross-linked peptides were loaded onto a picochip column (C18, 3 μm particle size, 300 Å pore size, 50 μm × 10.5 cm; New Objective) and eluted using a 60 min LC gradient: 5% B–8% B, 0 – 5 min; 8% B – 32% B, 5 – 45 min; 32% B–100% B, 45 – 49 min; 100% B, 49 - 54 min; 100% B - 5 % B, 54 min - 54 min 10 sec; 5% B, 54 min 10 sec - 60 min 10 sec; mobile phase A consisted of 0.1% formic acid (FA), and mobile phase B consisted of 0.1% FA in 80% acetonitrile. The QE HF-X instrument was operated in the data-dependent mode, where the top 8 most abundant ions (mass range 380–2,000, charge state 3 - 7) were fragmented by high-energy collisional dissociation (normalized collision energy 27). The target resolution was 120,000 for MS and 15,000 for MS/MS analyses. The quadrupole isolation window was 1.8 Th and the maximum injection time for MS/MS was set at 120 ms. After MS analysis, the data was searched by pLink for the identification of cross-linked peptides. The mass accuracy was specified as 10 and 20 p.p.m. for MS and MS/MS, respectively. Other search parameters included cysteine carbamidomethylation as a fixed modification and methionine oxidation as a variable modification. A maximum of three trypsin missed-cleavage sites was allowed. The initial search results were obtained using the default 5% false discovery rate, estimated using a target-decoy search strategy. The crosslink spectra were then manually checked to remove potential false-positive identifications essentially as previously described (Shi et al., 2014; Shi et al., 2015; Xiang et al., 2020).

### Site-directed mutagenesis

Mammalian expression plasmid of HSA was obtained from Addgene. E400R point mutation was introduced to the HSA sequence by the Q5 site-directed mutagenesis kit using the primer HSA-F and HSA-R. After sequence verification by Sanger Sequencing, plasmids bearing wild type HSA and the mutant were transfected to HeLa cells using Lipofectamine 3000 transfection kit (Invitrogen) and Opti-MEM (Gibco) according to the manufacturer's protocol. The cells were cultured overnight before change of medium to DMEM without FBS supplements to remove BSA. After a 48 h culture at 37°C, 5% CO_2_, the media expressing HSA were collected and stored at −20°C. The media were analyzed by SDS-PAGE and Western Blotting to confirm protein expression.

The PDZ domain (in the pGEX6p-1 vector) was obtained from the General Biosystems. A double point mutant of PDZ (i.e., R46E: K48D) was introduced by the Q5 Site-directed mutagenesis kit using specific primers of PDZ-F and PDZ-R. After verification by Sanger Sequencing, the mutant vector was transformed into BL21(DE3) cells (Thermo Scientific) for expression. The GST fusion PDZ mutant protein was purified by GSH resin as previously described.

### Fluorescence Microscopy

COS-7 cells were plated onto the glass bottom dish at an initial confluence of 60-70% and cultured overnight to let the cells attach to the dish. Cells were stained with MitoTracker Orange CMTMRos (1:4000) at 37 °C for 30 minutes, washed once with PBS and fixed with pre-cold methanol/ethanol (1:1) for 10 minutes. After being washed with PBS, the cells were blocked with 5% BSA for 1 hour. Alexa Fluor™ 647-conjugated Nb (1:100) was then added to the cells, incubated for 15 minutes at room temperature. Two-color wide-field fluorescence images were acquired using our custom-built system on an Olympus IX71 inverted microscope frame with 561 nm and 642 nm excitation lasers (MPB Communications, Pointe-Claire, Quebec, Canada) and a 100X oil immersion objective (NA=1.4, UPLSAPO 100XO; Olympus).

### The cleavage rules for in-silico digestion of Nbs by different proteases

Trypsin: C-terminal to K/R, not followed by P
Chymotrypsin: C-terminal to W/F/L/Y, not followed by P
GluC: C-terminal to D/E, not followed by P
AspN: N-terminal to D
LysC: C-terminal to K

### Sequence alignment of Nb database

Nb sequences were numbered using the software ANARCI according to Martin numbering scheme(MacCallum et al., 1996). Three CDRs (CDR1-CDR3) and four Framework sequences (FR1-FR4) were colored based on AbM definition. Alignments below the threshold e-value of 100 and introduced gaps with frequency of more than 0.999 were removed and the remaining sequences were plotted by WebLogo in **Figure 1A**.

### In-silico digestion of Nb database by different proteases and analysis of Nb CDR3 mapping

A high-quality database containing approximately 0.5 million unique Nb sequences was *in-silico* digested using different enzymes including trypsin,chymotrypsin, LysC, GluC, and AspN according to the above cleavage rules. CDR3 containing peptides were obtained to calculate the sequence coverages. The CDR3 coverages were then summed to generate **Figure 1D**,**Figure S1B** and **S1C**. The CDR3 peptide length distributions (by trypsin and chymotrypsin) were plotted to generate **Fig 1E**.

### Simulation of trypsin and chymotrypsin-aided MS mapping of Nbs

10,000 Nb sequences with unique CDR3 fingerprint sequences were randomly selected from the database. The selected Nbs were then *in-silico* digested by either trypsin or chymotrypsin (with no-miscleavage sites allowed) to generate CDR3 peptides. The following criteria were applied to these peptides to better simulate Nb identifications by MS: 1) peptides of favorable sizes for bottom-up proteomics (between 850- 3,000 Da) were first selected. 2) Peptides containing the highly conserved C-terminal FR4 motif of WGQGQVTS were further discarded. Based on our observations, such peptides are often dominated by C terminal y ion fragmentations, while having poorly fragmented ions on the CDR3 sequence which are essential for unambiguous CDR3 peptide identifications. 3) CDR3 peptides with limited Nb fingerprint information (containing less than 30% CDR3 sequence coverage) were removed. As a result, 2,111 unique tryptic peptides and 5,154 unique chymotryptic peptides were obtained. These peptides were then used to map Nb proteins. After protein assembly, only Nb identifications with sufficiently high CDR3 fingerprint sequence coverages (≥ 60%) were used to generate the venn diagram in **Fig 1F**.

### Phylogenetic analysis of Nb CDR3 sequences

Phylogenetic trees were generated by Clustal Omega with the input of unique Nb CDR3 sequences and the additional flanking sequences (i.e., YYCAA to the N-term and WGQG to the C-term of CDR3 sequences) to assist alignments. The data was plotted by ITol (Interactive Tree of Life). Isoelectric points and hydrophobicities of Nb CDR3s were calculated using the BioPython library.

### Evaluation of the reproducibility of Nb peptide quantification

Shared peptide identifications among different LC runs were used to evaluate the reproducibility of the label-free quantification method. For a typical 90 min LC gradient, the peptide peak width or full width at half maximum (FWHM) in general was less than 5s. The differences of peptide retention time among different LC runs were calculated to generate the kernel density estimation plots in **Fig 3B**. Peptide retention times from different LC runs were used to calculate pearson correlation and were plotted in **Figure S3B**.

### Sequence alignment and analysis of HSA and Llama serum albumin

Llama (Camelus Ferus) serum albumin sequence was fetched and aligned with HSA by tblastn (NCBI). The isoelectric point(pI) and hydropathy values for individual amino acids were obtained online from (https://www.peptide2.com/N_peptide_hydrophobicity_hydrophilicity.php). These values were normalized between 0 to 1.0 and the sequence variations between the two albumins were calculated for each aligned position (the pairwise differences of pI and hydropathy). For a specific aligned residue position, a value of 0 indicates identical residues were found between the two sequences, while 1.0 indicates the largest sequence variation, such as a charge reversion from the negatively charged residue glutamic acid 400 for HSA to the positively charged residue arginine at the corresponding aligned position for camelid albumin. A value of 0.5 was assigned at the position where an insertion or deletion of amino acid was identified. Sequence variations of both pI and hydropathy between HSA and Llama serum albumin were thus plotted. The plots were further smoothed by a gaussian function to generate **Fig 7B**.

### Analysis of relative abundance of amino acids on Nb CDRs

The amino acid frequencies at each CDR (including CDR1, CDR2 and CDR3 head) were calculated and normalized to generate the bar plots and the pie plots in **Figs 4, 7, S5 and S7**. CDR3 head sequences were obtained by removing the semi-conserved C terminal four residues of CDR3s. The CDR residue frequencies of both high-affinity and low-affinity Nbs were normalized based on the sum of the CDR residues of each affinity group.

### Analysis of amino acid positions on CDR3 heads

The relative position of a residue on a CDR3 head was calculated where a value of 0 indicates the very N terminus of a CDR3 head while 1.0 indicates the last residue. The CDR3 head sequences were then sliced into 20 bins with a bin width of 0.05. Within each bin, the occurrence of a specific type of amino acid (such as tyrosine, glycine, or serine) was counted and normalized to the sum of residues on CDR3 heads. The distributions of different amino acids including their relative positions and abundances were plotted in **Fig 4G and Fig S5E.**

### Proteomics database search of Nb peptide candidates

Raw MS data was searched by Sequest HT embedded in the Proteome Discoverer 2.1 against an in-house generated Nb sequence database using the standard target-decoy strategy for FDR estimation. The mass accuracy was specified as 10 ppm and 0.02 Da for MS1 and MS2, respectively. Other search parameters included cysteine carbamidomethylation as a fixed modification and methionine oxidation as a variable modification. A maximum of one or two missed-cleavage sites was allowed for trypsin and chymotrypsin-processed samples respectively. The initial search results were filtered by percolator with the FDR of 0.01 (strict) based on the q-value (Kall et al., 2007). After database search, the peptide-spectrum matches(PSMs) were exported, processed and analyzed by Augur Llama with following steps:

#### a. Nanobody Identification

##### i) Quality assessment of CDR3 fingerprints

Peptide candidates were first annotated as either CDR or FR peptides. To confidently identify CDR3 fingerprint peptides, we implemented a filter/algorithm requiring sufficient coverage of high-resolution CDR3 fragment ions in the PSMs(See illustration in **Fig S2B**). The filter was evaluated using a target sequence database containing approximately 0.5 million unique Nb sequences and a non-overlapping decoy database of similar size. Target and decoy Nb sequence databases herein used were obtained from different llamas. Any peptide identification from the decoy database was considered as a false positive. The FDR was defined based on the % of peptide identifications from the decoy database compared with those from the target database. CDR3 length was also considered to enable development of a sensitive CDR3 peptide filter. The CDR3 fragmentation coverage was defined as the percentage of the CDR3 residues that were matched by fragment ions (either b ions or y ions) within the mass accuracy window. Spectra of the same peptide were combined for assessment. Only CDR3 peptides that passed this filter (5% FDR) were selected for the downstream Nb assembly.

##### ii) Nanobody sequence assembly

CDR peptides including the confident CDR3 peptides were used for Nb protein assemblies. Two additional criteria must be matched before a Nb could be identified. These include: 1) both CDR1 and CDR2 peptides must be available for a Nb assembly. 2) for any Nb identification, a minimum of 50% combined CDR coverage was mandated.

#### b. Quantification and classification of antigen-specific Nb repertoires

MS raw data was accessed by MSFileReader 3.1 SP4(ThermoFisher), and a python library of pymsfilereader (https://github.com/frallain/pymsfilereader). Reliable CDR3 peptides that passed the quality filter were quantified by label-free LC/MS.

#### i) CDR3 peptide quantification

To enable accurate label-free quantification of a CDR3 peptide identification across different LC runs, different retention time windows for peptide peak extraction were specified. For peptides that could be directly identified by the search engine based on the MS/MS spectra, a small quantification window of +/− 0.5 minutes retention time (RT) shift was used for peak extractions. For peptides that were not directly identified from a particular LC run (due to the complexity of peptides and stochastic ion sampling), their RTs were predicted based on the RT of the adjacent LC and were adjusted using the median RT difference of the commonly identified peptides between the two LC runs. In this case, a relaxed RT window of +/− 2.0 minutes (for a typical 90 min LC gradient), in which approximately 95% of all the identified peptides can be matched between the two LC runs, was applied to facilitate extraction of the peptide peaks. Both m/z and z of a peptide were used for peak extractions with a mass accuracy window of +/− 10 ppm. The peptide peaks were extracted and smoothed using a Gaussian function. Their AUCs (area under the curve) were calculated and AUCs from the replicated LC runs were averaged to infer the CDR3 peptide intensities.

#### ii) Classifications of Nbs

To enable accurate classifications e.g., based on Nb affinities, relative ion intensities (AUCs) of the CDR3 fingerprint peptides among three different biochemically fractionated Nb samples (*F1*, *F2* and *F3*) were quantified as *I1*, *I2* and *I3*. Based on the quantification results, CDR3 peptides were arbitrarily classified into three clusters (*C1*, *C2*, and *C3*) using the following criteria:

1. For *C3* (high-affinity) cluster: *I3 > I1+I2* (indicating Nbs were more specific to *F3*)
2. For *C2* (mediocre-affinity) cluster: *I2 > I1+I3* (indicating Nbs were more specific to *F2*)
3. For *C1* (low-affinity) cluster:
*I1> I2+I3* (indicating Nbs were either more specific to *F1* or likely nonspecific binders), alternatively, if *I1< I2+I3* and *I2< I1+I3* and *I3< I1+I2,* these Nb identifications were likely nonspecifically identified and were grouped into *C1* as well. See illustration in **Fig S2C.**

The above method was used to classify HSA and GST Nbs. Some modifications were made for quantification and characterization of high-affinity PDZ Nbs. Specifically, an additional control of MBP interacting Nbs “ *F_control*” (ion intensity of *I_control*) was included for quantification. High-affinity cluster Nbs (represented by their unique CDR3 peptides) were defined when the sum intensities of *I2* and *I3* for a Nb CDR3 peptide were 20 fold higher than *I_control*(i.e. *20*I_control < I2+I3*). For Nbs where more than one unique CDR3 peptides were used for quantification, classification results among different CDR3 peptides from the same Nb must be consistent; otherwise they were removed before the final results were reported.

### Heatmap analysis of the relative intensities of CDR3 peptides

The identified CDR3 peptides were quantified based on their relative MS1 ion intensities and were subsequently clustered using scripts in Augur Llama. Z-scores were calculated based on the relative ion intensities and were used to generate a heatmap in **Fig 3A** for visualization.

### Structural modeling of antigen-Nb complexes

Structural models for Nbs were obtained using a multi-template comparative modeling protocol of MODELLER(Webb and Sali, 2014). Next, we refine the CDR3 loop and select the top 5 scoring loop conformations for the downstream docking. Each Nb model is then docked to the respective antigen by an antibody-antigen docking protocol of PatchDock software that focuses the search to the CDRs (Schneidman-Duhovny et al., 2005). The models are then re-scored by a statistical potential SOAP (Dong et al., 2013). The antigen interface residues (distance <6Å from Nb atoms) among the 10 best scoring models according to the SOAP score were used to determine the epitopes. Once the epitopes were defined, we clustered Nbs based on the epitope similarity using k-means clustering. The clusters reveal the most immunogenic surface patches on the antigens. Antigen-Nb complexes with CXMS data were modeled by distance-restrained based PatchDock protocol that optimizes restraints satisfaction (Schneidman-Duhovny and Wolfson, 2020). A restraint was considered satisfied if the Ca-Ca distance between the cross-linked residues was within 25Å and 20Å for DSS and EDC cross-linkers, respectively (Fernandez-Martinez et al., 2016; Shi et al., 2014). In the case of ambiguous restraints, such as the GST dimer, we require that one of the possible cross-links is satisfied.

### Measurement of the interface curvature

The interface curvature was calculated as the average of the shape function of the interface atoms of the antigen or the Nb. For this purpose a sphere of radius R (6Å) is placed at a surface point of the interface atom. The fraction of the sphere inside the solvent-excluded volume of the protein is the shape function at the corresponding atom (Connolly, 1986).

### Machine learning analysis of Nb repertoires

We trained a deep neural network to distinguish between low- and high- affinity Nbs that were characterized by the accurate high-pH fractionation method and quantitative proteomics. Our model consists of one convolutional layer with batch normalization and ReLU activation function, followed by a max pooling layer ending with a fully connected layer to integrate the features extracted into the logits layer that leads to the classifier prediction. The convolutional layer consists of 20 1D filters, representing local receptive fields with window size of 7 amino acids, long enough to capture the relevant CDRs and short enough to avoid data overfitting. During the forward pass, each filter slides along the protein sequence with a fixed stride performing an elementwise multiplication with the current sequence window, followed by summing it up to generate a filter response. The classification accuracy of the model was 92%.

To understand the physicochemical features learned by the network for distinguishing low- and high- affinity binders we calculated the activation path through the network back from the prediction to the activated filter. Similarly to the backpropagation algorithm, we iterated backward from the last two layers of fully connected network, extracting for each sequence the output signal and looking for the highest peaks which contribute the most weight to the classification. In the same way, we calculated upstream the contribution of each filter to those peaks. In addition, we analyzed filter activity in CDRs to extract region-specific dominant filters. This process of network interpretation results in a unique contribution per filter per sequence. Each filter is activated along the sequence downsampled in the max pooling layer. For each filter, we then picked its highest peak leading to classification. Finally we looked at the most contributing filters per sequence and there also we got an interesting filter out with more than 30% contribution in those regions of interest.

## Acknowledgements

We thank J.L.Jia and Y.Jiao for the NGS experiments, Z.Shen and Z.Y.Xiao for assistance in the mutagenesis experiments, J.W. Ahn and P. Duplex for the help on the biophysic experiments. J.J.Wang (Rockefeller University) for providing a script for the preliminary database analysis. Y.Liu, H.Maayan and D.Groebe for critical comments and assistance. This work was supported by University of Pittsburgh School of Medicine (Y.S.) and a UPMC Aging Institute pilot fund (Y.S.), ISF 1466/18, Israel ministry of Science and Technology and HUJI-CIDR (D.S.).

## Contributions

Y.S. conceived the research. Y.S. and D.S. supervised the study. Y.X. performed all the wet experiments. Y.X. and Y.S. performed the MS analysis. Z.S., D.S., Y.S., Y.X. and L.B. developed the informatics tools and analyzed data. J.X and Y.L facilitated the fluorescence imaging experiments. Y.S. drafted the manuscript. Y.S., D.S., X.Y., Z.S. edited the manuscript.

## Completing Interests

The University of Pittsburgh has filed a provisional patent in connection to the manuscript.

---

**Figure S1.**
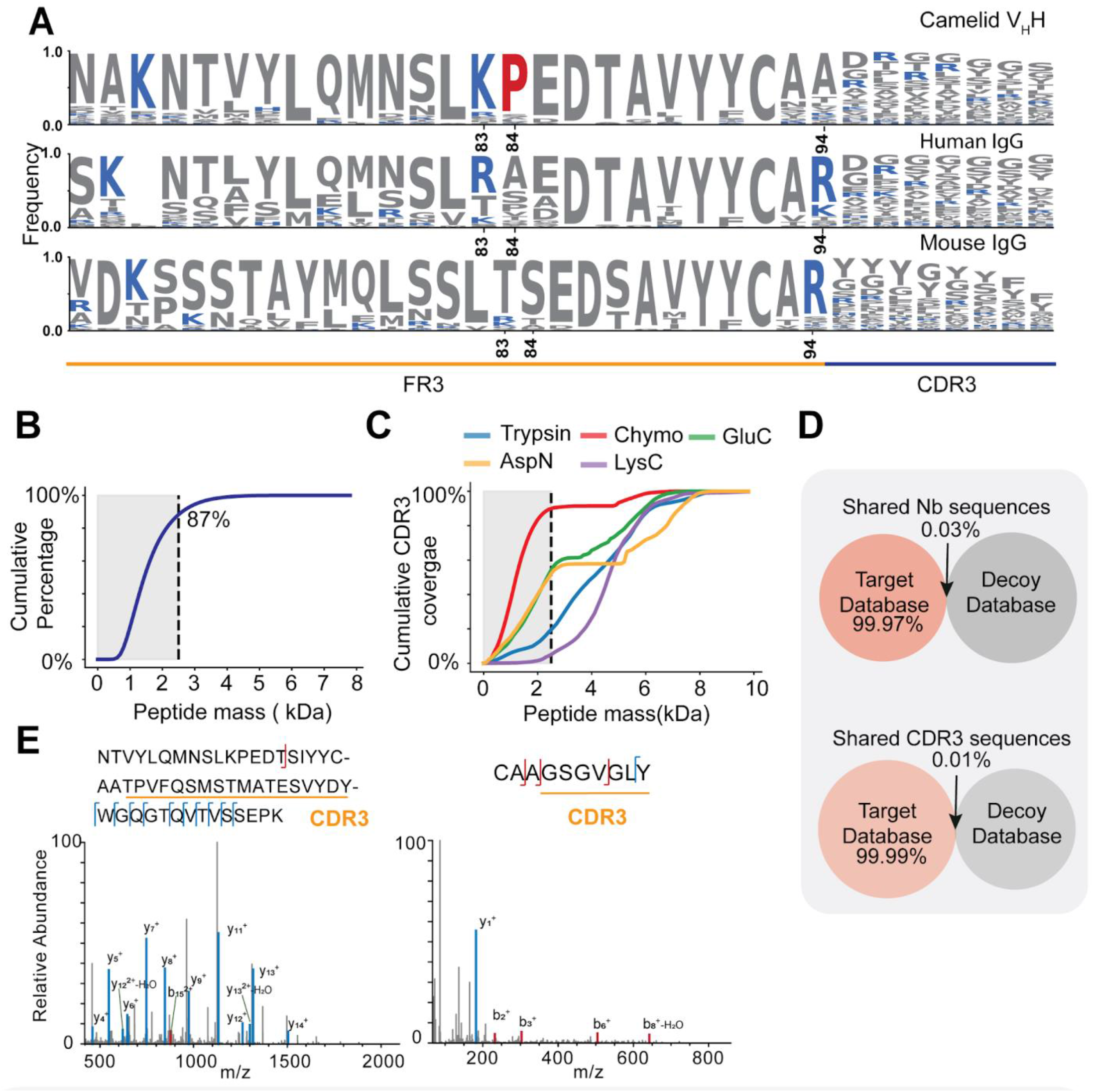
Analysis of NGS Nb databases and representative false positive CDR3 peptide identifications (Related to Figure 1 and Methods). **A.** Sequences logos comparison among camelid VHH, human lgG and Mouse lgG shows different sequence features. Only FR3 and partial CDR3 were displayed. Lysine and arginine (Trypsin putative cleavage sites) were colored in blue. Boxed area shows more conserved praline found in camelid V_H_H. **B.** The mass distribution of ~1.5 million peptide identifications of human proteins from PeptideAtlas. **C.** *In silico* digestion of Nb NGS database by different proteases (AspN, GluC, LysC, Trypsin and Chymotrypsin) and plot of peptide masses. **D.** The overlaps between the target Nb sequence database of the immunized Llama and a decoy database from another native Llama. ~ 0.5 million sequences were included in each database. **E.** A representative low quality/false positive MS/MS spectrum (HCD) of a tryptic CDR3 peptide. **F.** That of a chymotryptic CDR3 peptide. Few high-resoluton fragment ions were matched in the spectra.

**Figure S2.**
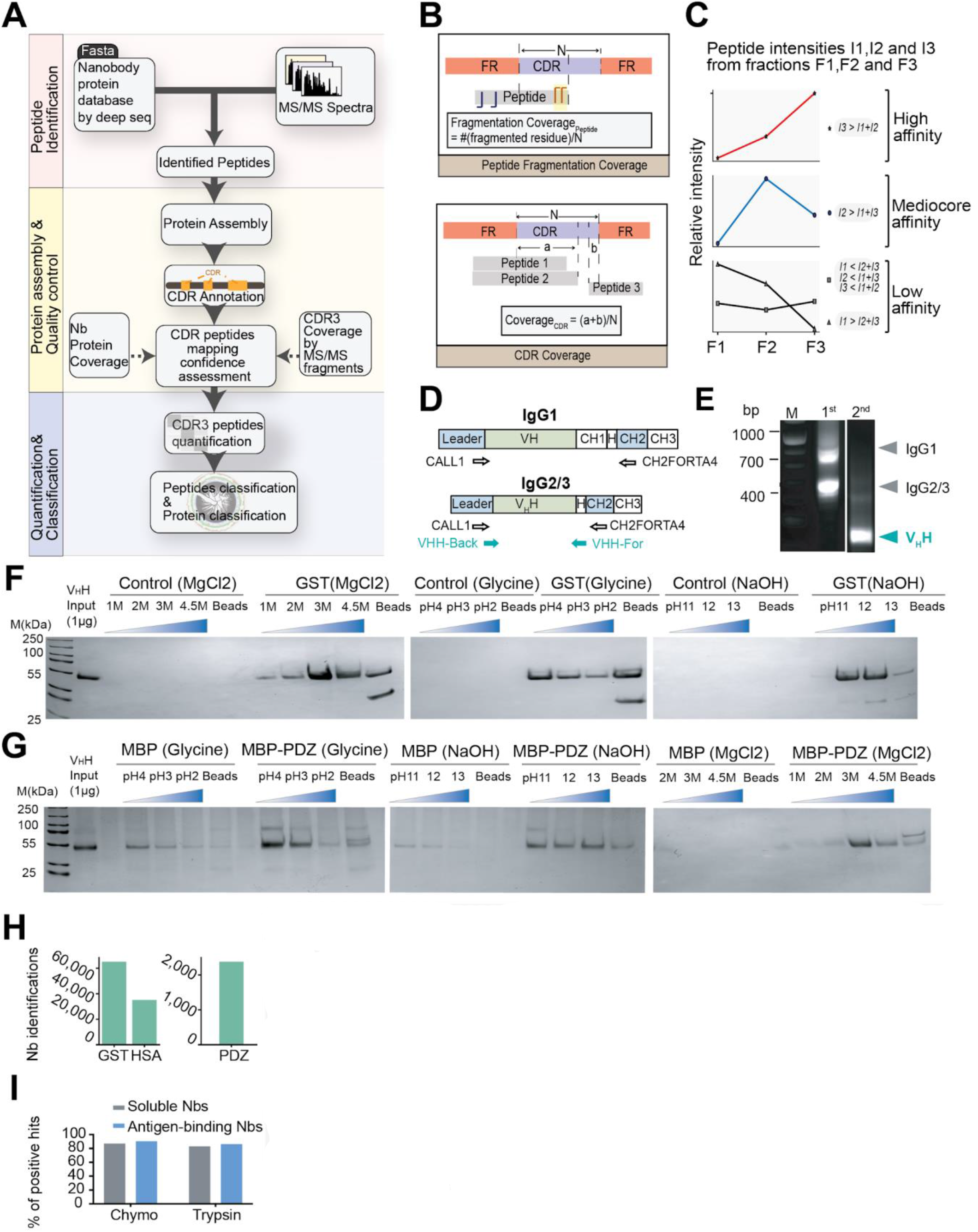
The informatics pipeline of “Augur Llama” for Nb proteomics and validation of Nb binders (Related to Figure 2 and Methods). **A.** Schematics of the informatic pipeline. Three modules including 1) peptide identifications, 2) Nb peptide and protein quality control, and 3) quantification and classifications were presented. Nb proteomics data is first searched against the search engine. The initial identifications that pass the search engine will be automatically annotated, and evaluated based on different quality filters at peptide and protein levels. High-quality fingerprint peptides that pass the quality filters will be quantified and clustered. **B.** Illustrations of the Nb CDR3 spectrum and coverage quality filters. **C.** lllusreations of peptide classification method. **D.** Schematic of PCR amplifications of HcAb variable domain (VHH) from B lympcytes of the camel id. **E.** DNA gel electrophoresis of the VHH PCR amplicons from the cDNA libraries prepared from the immunized bone marrow/blood. **F.** SOS-PAGE analysis of fractionated GST-Nbs based on different fractionation protocols. **G.** SOS-PAGE analysis of PDZ Nbs. Maltose-binding protein (MBP) tag was fused to PDZ domain and the fusion protein was used as affinity handle for isolation. MBP was used as a negative control for quantification. **H.** Unique Nb identifications for different antigens. **I.** Comparison of antigen-specific Nbs identified by either chymotrypsin or trypsin-based method. Y axis stands for the % of the positive hits that were randomly selected for verifications.

**Figure S3.**
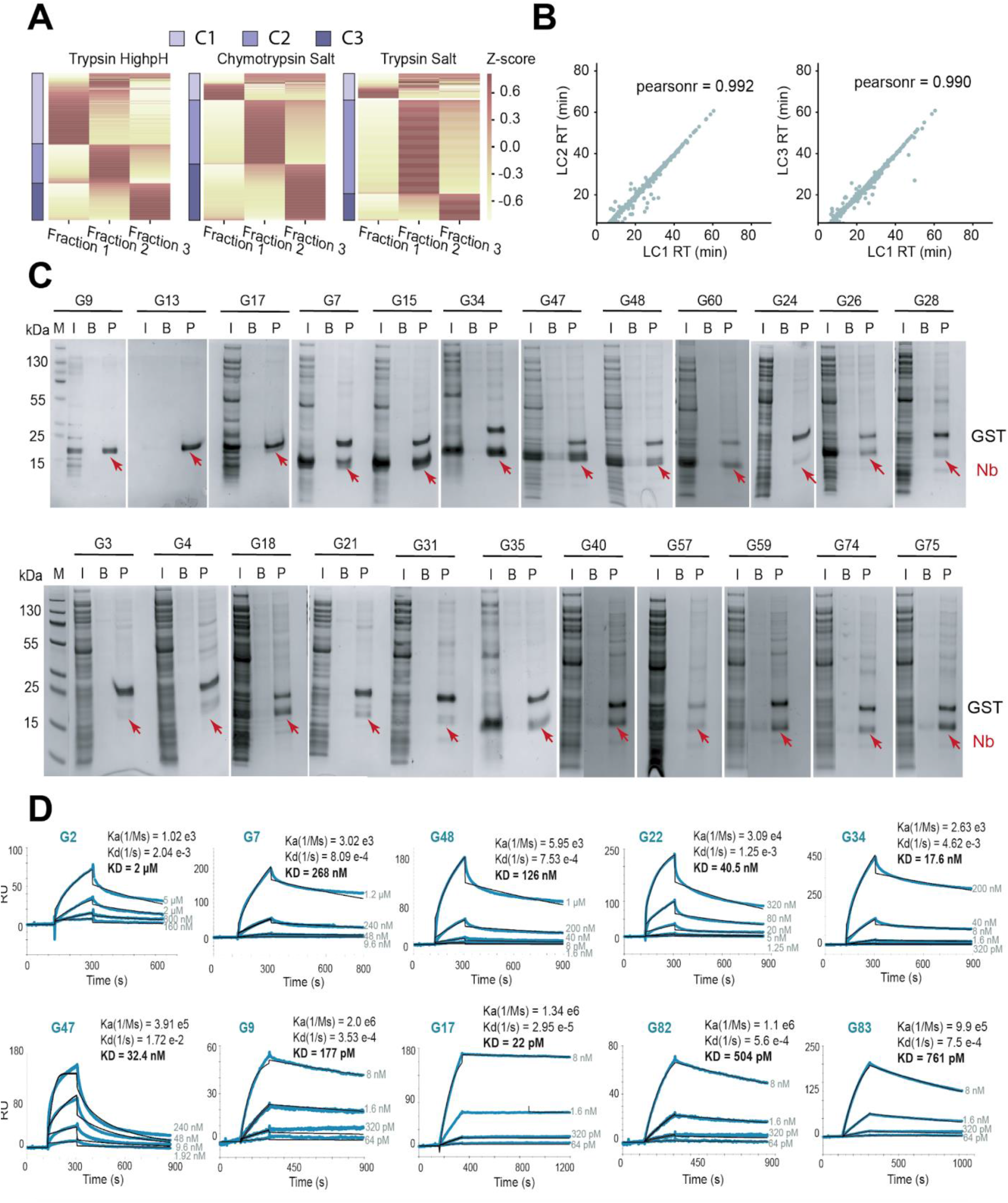
Proteomic quantifications, biochemical verifications and affinity measurements of Nb_GST_ (Related to Figure 3 and Method). **A.** Proteomic quantifications and heatmap analysis of Nb_GST_ based on different fractionation methods. **B.** Pearson correlations of LC retention times of different fractionated Nb peptide samples. **C.** Representative GST beads-binding assay. GST coupled resin was used to specifically isolate recombinant Nb from the *E.coli* lysis. Red arrows indicate enriched Nbs. Inactivated resin was used for negative control. **D.** SPR kinetic measurements of 10 representative Nb_GST_

**Figure S4.**
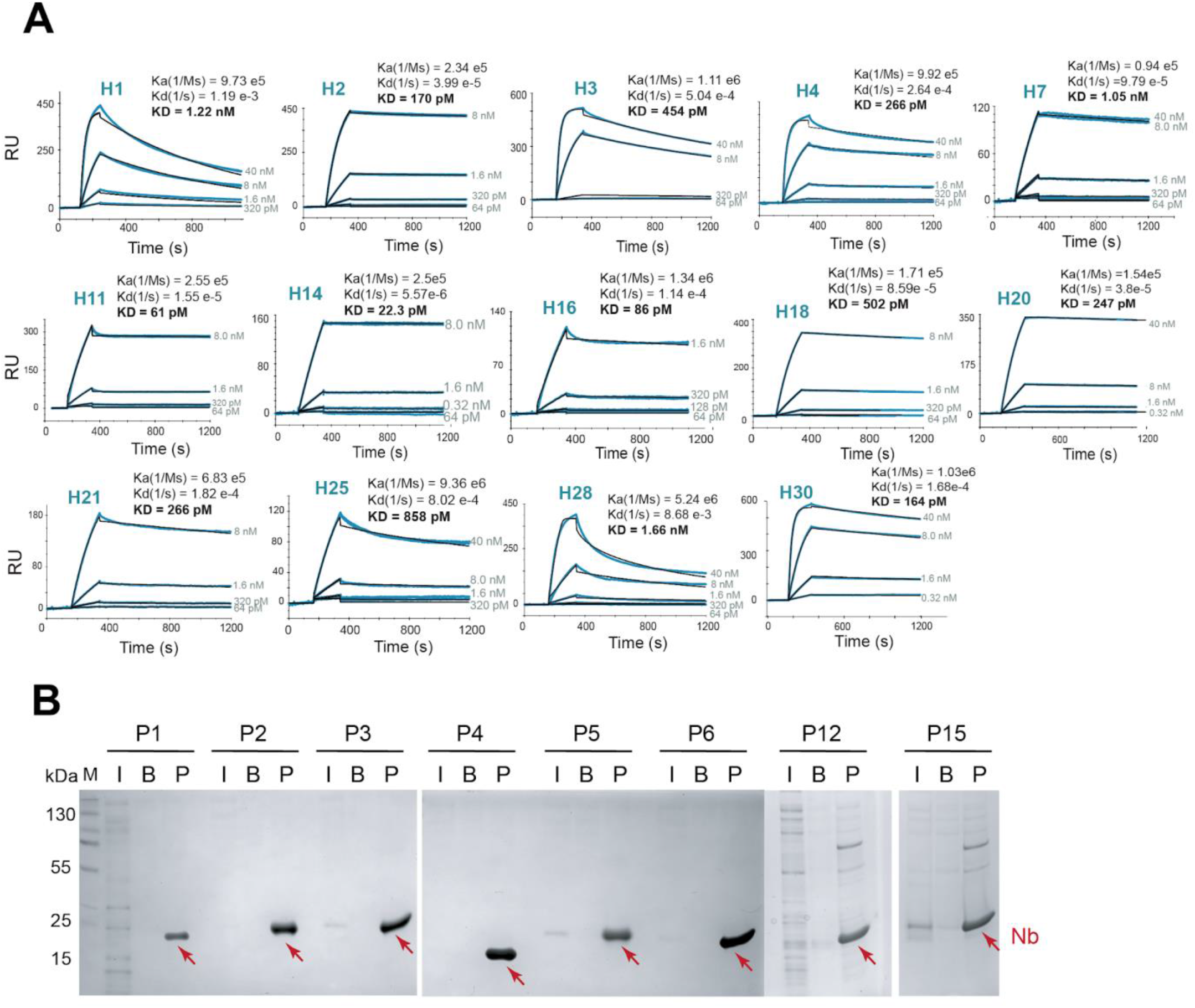
Characterizations of High-quality HSA and PDZ Nbs (Related to Figure 3 and Methods). **A.** SPR kinetic measurements of representative high-affinity Nb_HSA_. **B.** Beads-binding assays of selected high-quality Nb_PDZ_. Recombinant MBP fusion PDZ was used as affinity handle for isolation of Nbs from *E.coli* lysates. MBP coupled resin was used for negative control. I: *E.coli* lysate input, B: beads control, P: affinity pullout by PDZ.

**Figrue S5.**
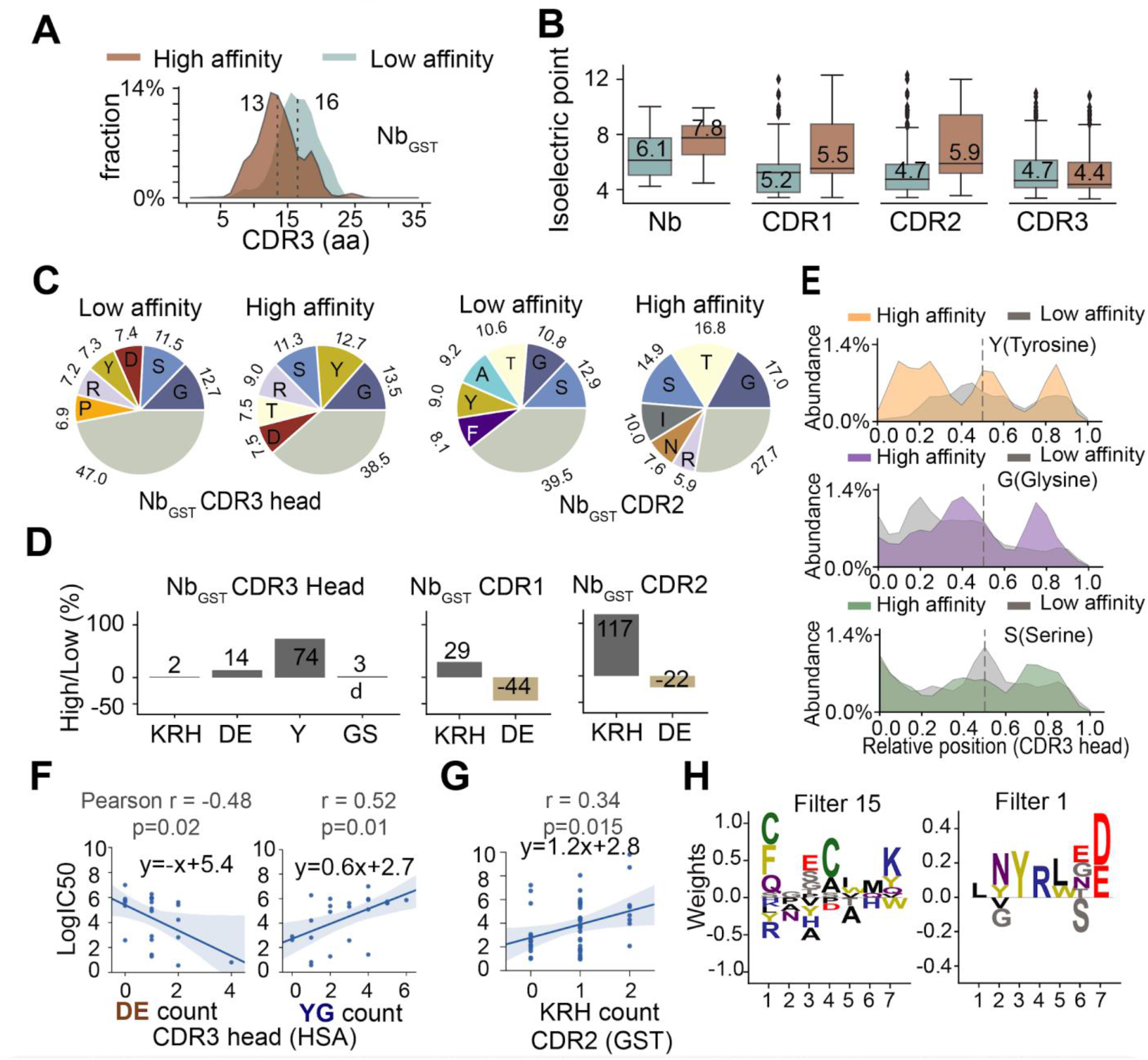
The mechanisms of NbGsr antibody affinity maturation (Related to Figure 4 and Methods). **A.** Distributions of CDR3 lengths of high-affinity (brown) and low-affinity (steel) Nb_GST_ **B.** Comparisons of pis of low- and high-affinity Nb_GST_ and contri bution by CDRs_GST_ **C.** Pie charts of the amino acid compositions of the CDR3_GST_ heads and the CDR2_GST_ Only the top 6 abundant residues are shown. **D.** The relative changes of abundant amino acids on CDR3_GST_ heads. Positive charged residues of K (lysine)/ R (arginine)/ H (histidine), negative charged residues of D (aspartic acid)/E (glutamic acid), aromatic residue ofY(tyrosine) and small flexible amino acids of G (glycine)/ S (serine) are shown. **E.** Comparison of the relative position of Y (tyrosine), G (glycine) and S (serineS) on the CDR3_GST_ heads. **F.** Correlation plots of the ELISA affinities and the number of specific amino acids on the CDR3HsA heads. Pearson correlation coefficients and the statistical values are shown. **G.** The correlation plot of ELISA affinities and the number of positively charged residues on the CDR2_GST_ **H.** Sequence logo of two representative convolutional CDR3 filters (Filter 15 for low-affinity Nb_GST_; filter 1 for high-affinity Nb_GST_l learned by a deep learning model.

**Figure S6.**
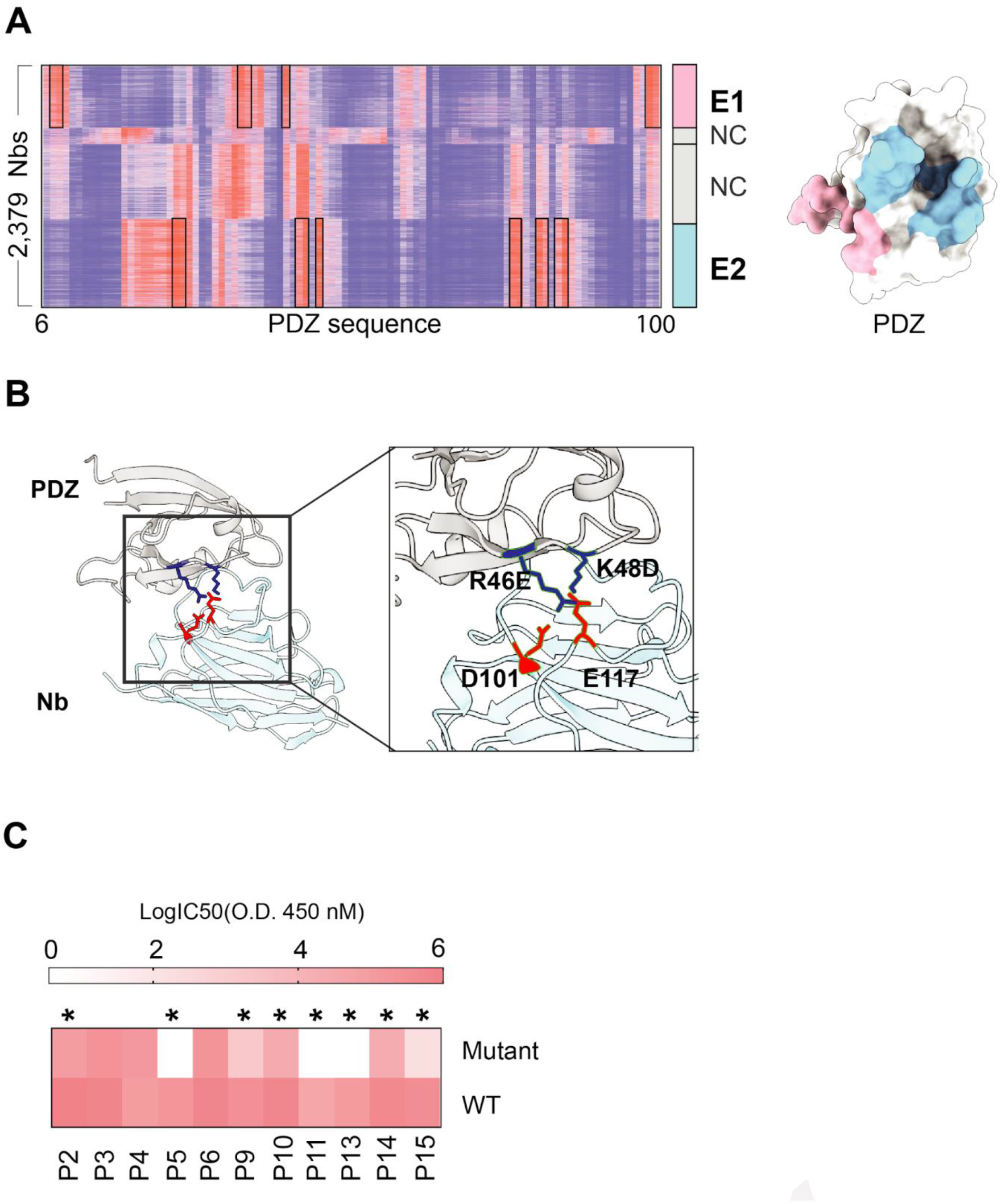
High-throughput docking and mutagenesis analysis of PDZ-Nb complexes (Related to Figure 5 and 6). **A.** Heatmap analysis of structural docking of 2,370 different PDZ-Nb complexes showing two converged epitopes (E 1: 75-88, 143-148; E2: 33-43). Surface representations of the PDZ epitopes were shown. E1 and E2 were in pink and sky blue, respectively. **B.** Structural model of a representative PDZ-Nb complex showing the formation of two salt bridges on the E2 interface. **C.** ELISA affinity screening (heatmap) of 11 different Nbs for binding to PDZ and the point mutant (R46E:K48O). * indicates decreased affinity.

**Figure S7.**
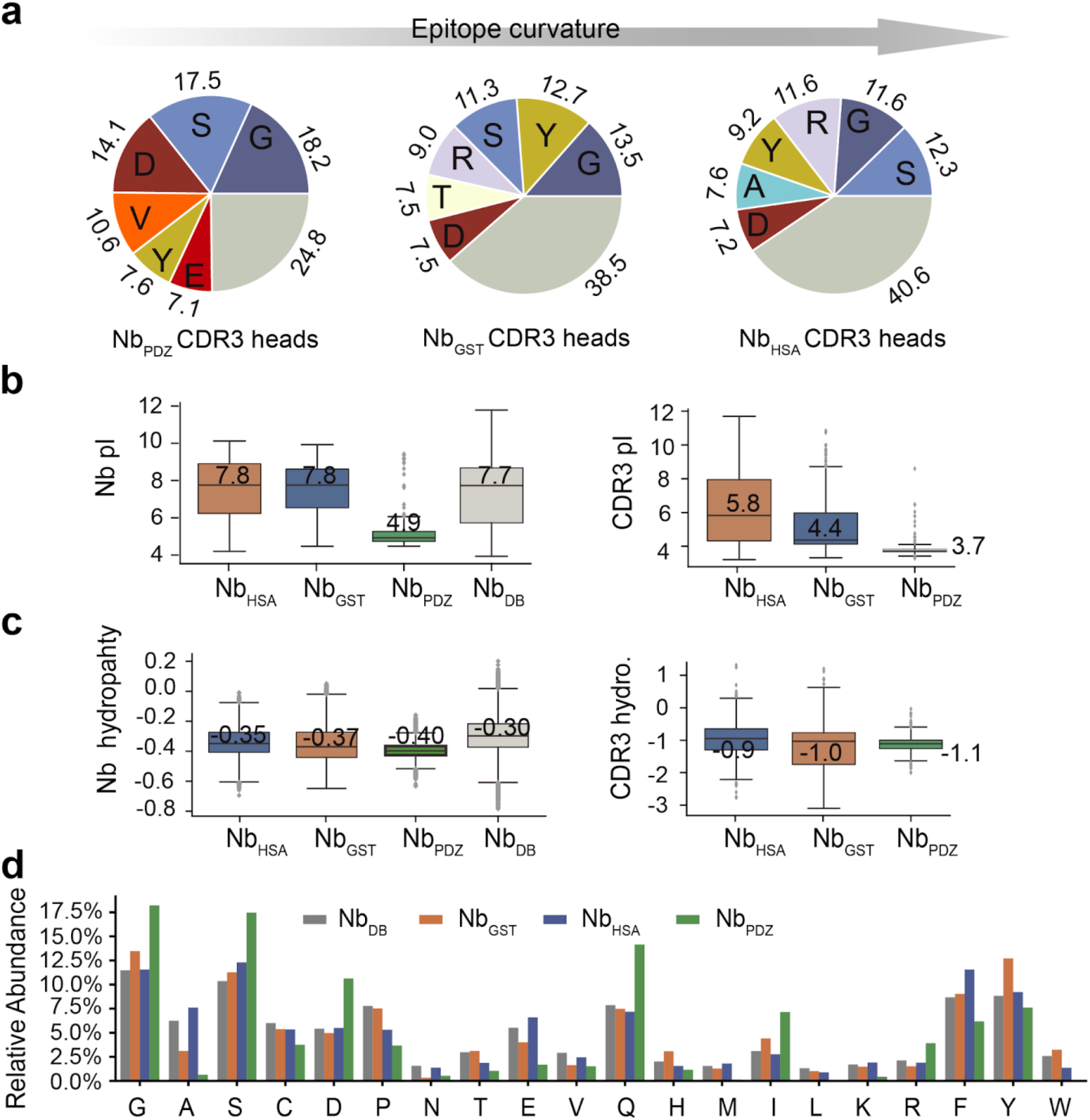
Comparison among different antigen-specific Nbs (Related to Figure 7). **A**. Pie charts of the top 6 most abundant amino acids on the Nb CDR3 heads. **B**. Comparisons of pl among different Nbs. **C**. Comparisons of hydropathy among different Nbs. **D**. Relative abundance of different amino acids on the CDR3 heads. DB: NGS Nbs sequence database.

## Reference

Abhinandan, K.R., and Martin, A.C. (2008). Analysis and improvements to Kabat and structurally correct numbering of antibody variable domains. Mol Immunol 45, 3832–3839.

Akram, A., and Inman, R.D. (2012). Immunodominance: a pivotal principle in host response to viral infections. Clin Immunol 143, 99–115.

Arbabi Ghahroudi, M., Desmyter, A., Wyns, L., Hamers, R., and Muyldermans, S. (1997). Selection and identification of single domain antibody fragments from camel heavy-chain antibodies. FEBS Lett 414, 521–526.

Baran, D., Pszolla, M.G., Lapidoth, G.D., Norn, C., Dym, O., Unger, T., Albeck, S., Tyka, M.D., and Fleishman, S.J. (2017). Principles for computational design of binding antibodies. Proc Natl Acad Sci U S A 114, 10900–10905.

Beghein, E., and Gettemans, J. (2017). Nanobody Technology: A Versatile Toolkit for Microscopic Imaging, Protein-Protein Interaction Analysis, and Protein Function Exploration. Front Immunol 8, 771.

Chait, B.T., Cadene, M., Olinares, P.D., Rout, M.P., and Shi, Y. (2016). Revealing Higher Order Protein Structure Using Mass Spectrometry. J Am Soc Mass Spectrom 27, 952–965.

Chaplin, D.D. (2010). Overview of the immune response. J Allergy Clin Immunol 125, S3–23.

Chen, Z.L., Meng, J.M., Cao, Y., Yin, J.L., Fang, R.Q., Fan, S.B., Liu, C., Zeng, W.F., Ding, Y.H., Tan, D., et al. (2019). A high-speed search engine pLink 2 with systematic evaluation for proteome-scale identification of cross-linked peptides. Nat Commun 10, 3404.

Cheung, W.C., Beausoleil, S.A., Zhang, X., Sato, S., Schieferl, S.M., Wieler, J.S., Beaudet, J.G., Ramenani, R.K., Popova, L., Comb, M.J.*, et al.* (2012). A proteomics approach for the identification and cloning of monoclonal antibodies from serum. Nat Biotechnol 30, 447–452.

Chevalier, A., Silva, D.A., Rocklin, G.J., Hicks, D.R., Vergara, R., Murapa, P., Bernard, S.M., Zhang, L., Lam, K.H., Yao, G.*, et al.* (2017). Massively parallel de novo protein design for targeted therapeutics. Nature 550, 74–79.

Connolly, M.L. (1986). Shape complementarity at the hemoglobin alpha 1 beta 1 subunit interface. Biopolymers 25, 1229–1247.

Cox, J., and Mann, M. (2008). MaxQuant enables high peptide identification rates, individualized p.p.b.-range mass accuracies and proteome-wide protein quantification. Nat Biotechnol 26, 1367–1372.

Crooks, G.E., Hon, G., Chandonia, J.M., and Brenner, S.E. (2004). WebLogo: a sequence logo generator. Genome Res 14, 1188–1190.

Curry, S., Mandelkow, H., Brick, P., and Franks, N. (1998). Crystal structure of human serum albumin complexed with fatty acid reveals an asymmetric distribution of binding sites. Nat Struct Biol 5, 827–835.

DeKosky, B.J., Ippolito, G.C., Deschner, R.P., Lavinder, J.J., Wine, Y., Rawlings, B.M., Varadarajan, N., Giesecke, C., Dorner, T., Andrews, S.F., et al. (2013). High-throughput sequencing of the paired human immunoglobulin heavy and light chain repertoire. Nat Biotechnol 31, 166–169.

Desmyter, A., Transue, T.R., Ghahroudi, M.A., Thi, M.H., Poortmans, F., Hamers, R., Muyldermans, S., and Wyns, L. (1996). Crystal structure of a camel single-domain VH antibody fragment in complex with lysozyme. Nat Struct Biol 3, 803–811.

Dong, G.Q., Fan, H., Schneidman-Duhovny, D., Webb, B., and Sali, A. (2013). Optimized atomic statistical potentials: assessment of protein interfaces and loops. Bioinformatics 29, 3158–3166.

Doyle, D.A., Lee, A., Lewis, J., Kim, E., Sheng, M., and MacKinnon, R. (1996). Crystal structures of a complexed and peptide-free membrane protein-binding domain: molecular basis of peptide recognition by PDZ. Cell 85, 1067–1076.

Dunbar, J., and Deane, C.M. (2016). ANARCI: antigen receptor numbering and receptor classification. Bioinformatics 32, 298–300.

Egloff, P., Zimmermann, I., Arnold, F.M., Hutter, C.A.J., Morger, D., Opitz, L., Poveda, L., Keserue, H.A., Panse, C., Roschitzki, B., et al. (2019). Engineered peptide barcodes for in-depth analyses of binding protein libraries. Nat Methods 16, 421–428.

Elias, J.E., and Gygi, S.P. (2007). Target-decoy search strategy for increased confidence in large-scale protein identifications by mass spectrometry. Nat Methods 4, 207–214.

Els Conrath, K., Lauwereys, M., Wyns, L., and Muyldermans, S. (2001). Camel single-domain antibodies as modular building units in bispecific and bivalent antibody constructs. J Biol Chem 276, 7346–7350.

Fernandez-Martinez, J., Kim, S.J., Shi, Y., Upla, P., Pellarin, R., Gagnon, M., Chemmama, I.E., Wang, J., Nudelman, I., Zhang, W., et al. (2016). Structure and Function of the Nuclear Pore Complex Cytoplasmic mRNA Export Platform. Cell 167, 1215–1228 e1225.

Finn, J.A., Koehler Leman, J., Willis, J.R., Cisneros, A., 3rd, Crowe, J.E., Jr., and Meiler, J. (2016). Improving Loop Modeling of the Antibody Complementarity-Determining Region 3 Using Knowledge-Based Restraints. PLoS One 11, e0154811.

Fridy, P.C., Li, Y., Keegan, S., Thompson, M.K., Nudelman, I., Scheid, J.F., Oeffinger, M., Nussenzweig, M.C., Fenyo, D., Chait, B.T., et al. (2014). A robust pipeline for rapid production of versatile nanobody repertoires. Nat Methods 11, 1253–1260.

Hamers-Casterman, C., Atarhouch, T., Muyldermans, S., Robinson, G., Hamers, C., Songa, E.B., Bendahman, N., and Hamers, R. (1993). Naturally occurring antibodies devoid of light chains. Nature 363, 446–448.

Inbar, D., Hochman, J., and Givol, D. (1972). Localization of antibody-combining sites within the variable portions of heavy and light chains. Proc Natl Acad Sci U S A 69, 2659–2662.

Kall, L., Canterbury, J.D., Weston, J., Noble, W.S., and MacCoss, M.J. (2007). Semi-supervised learning for peptide identification from shotgun proteomics datasets. Nat Methods 4, 923–925.

Kim, S.J., Fernandez-Martinez, J., Nudelman, I., Shi, Y., Zhang, W., Raveh, B., Herricks, T., Slaughter, B.D., Hogan, J.A., Upla, P., et al. (2018). Integrative structure and functional anatomy of a nuclear pore complex. Nature 555, 475–482.

Lam, A.Y., Pardon, E., Korotkov, K.V., Hol, W.G.J., and Steyaert, J. (2009). Nanobody-aided structure determination of the EpsI:EpsJ pseudopilin heterodimer from Vibrio vulnificus. J Struct Biol 166, 8–15.

Lawrence, M.C., and Colman, P.M. (1993). Shape complementarity at protein/protein interfaces. J Mol Biol 234, 946–950.

Leitner, A., Faini, M., Stengel, F., and Aebersold, R. (2016). Crosslinking and Mass Spectrometry: An Integrated Technology to Understand the Structure and Function of Molecular Machines. Trends Biochem Sci 41, 20–32.

Letunic, I., and Bork, P. (2007). Interactive Tree Of Life (iTOL): an online tool for phylogenetic tree display and annotation. Bioinformatics 23, 127–128.

Li, T., Qi, S., Unger, M., Hou, Y.N., Deng, Q.W., Liu, J., Lam, C.M.C., Wang, X.W., Xin, D., Zhang, P., et al. (2016). Immuno-targeting the multifunctional CD38 using nanobody. Sci Rep 6, 27055.

MacCallum, R.M., Martin, A.C., and Thornton, J.M. (1996). Antibody-antigen interactions: contact analysis and binding site topography. J Mol Biol 262, 732–745.

McCoy, A.J., Chandana Epa, V., and Colman, P.M. (1997). Electrostatic complementarity at protein/protein interfaces. J Mol Biol 268, 570–584.

McMahon, C., Baier, A.S., Pascolutti, R., Wegrecki, M., Zheng, S., Ong, J.X., Erlandson, S.C., Hilger, D., Rasmussen, S.G.F., Ring, A.M., et al. (2018). Yeast surface display platform for rapid discovery of conformationally selective nanobodies. Nat Struct Mol Biol 25, 289–296.

Muyldermans, S. (2013). Nanobodies: natural single-domain antibodies. Annu Rev Biochem 82, 775–797.

Niethammer, M., Valtschanoff, J.G., Kapoor, T.M., Allison, D.W., Weinberg, R.J., Craig, A.M., and Sheng, M. (1998). CRIPT, a novel postsynaptic protein that binds to the third PDZ domain of PSD-95/SAP90. Neuron 20, 693–707.

Peng, H.P., Lee, K.H., Jian, J.W., and Yang, A.S. (2014). Origins of specificity and affinity in antibody-protein interactions. Proc Natl Acad Sci U S A 111, E2656–2665.

Pires, D.E., Ascher, D.B., and Blundell, T.L. (2014). mCSM: predicting the effects of mutations in proteins using graph-based signatures. Bioinformatics 30, 335–342.

Rini, J.M., Schulze-Gahmen, U., and Wilson, I.A. (1992). Structural evidence for induced fit as a mechanism for antibody-antigen recognition. Science 255, 959–965.

Rout, M.P., and Sali, A. (2019). Principles for Integrative Structural Biology Studies. Cell 177, 1384–1403.

Russel, D., Lasker, K., Webb, B., Velazquez-Muriel, J., Tjioe, E., Schneidman-Duhovny, D., Peterson, B., and Sali, A. (2012). Putting the pieces together: integrative modeling platform software for structure determination of macromolecular assemblies. PLoS Biol 10, e1001244.

Sanjuan, R., Nebot, M.R., Chirico, N., Mansky, L.M., and Belshaw, R. (2010). Viral mutation rates. J Virol 84, 9733–9748.

Scheid, J.F., Mouquet, H., Ueberheide, B., Diskin, R., Klein, F., Oliveira, T.Y., Pietzsch, J., Fenyo, D., Abadir, A., Velinzon, K., et al. (2011). Sequence and structural convergence of broad and potent HIV antibodies that mimic CD4 binding. Science 333, 1633–1637.

Schneidman-Duhovny, D., Inbar, Y., Nussinov, R., and Wolfson, H.J. (2005). PatchDock and SymmDock: servers for rigid and symmetric docking. Nucleic Acids Res 33, W363–367.

Schneidman-Duhovny, D., and Wolfson, H.J. (2020). Modeling of Multimolecular Complexes. Methods Mol Biol 2112, 163–174.

Sela-Culang, I., Kunik, V., and Ofran, Y. (2013). The structural basis of antibody-antigen recognition. Front Immunol 4, 302.

Sheehan, D., Meade, G., Foley, V.M., and Dowd, C.A. (2001). Structure, function and evolution of glutathione transferases: implications for classification of non-mammalian members of an ancient enzyme superfamily. Biochem J 360, 1–16.

Sheng, M., and Sala, C. (2001). PDZ domains and the organization of supramolecular complexes. Annu Rev Neurosci 24, 1–29.

Shi, Y., Fernandez-Martinez, J., Tjioe, E., Pellarin, R., Kim, S.J., Williams, R., Schneidman-Duhovny, D., Sali, A., Rout, M.P., and Chait, B.T. (2014). Structural characterization by cross-linking reveals the detailed architecture of a coatomer-related heptameric module from the nuclear pore complex. Mol Cell Proteomics 13, 2927–2943.

Shi, Y., Pellarin, R., Fridy, P.C., Fernandez-Martinez, J., Thompson, M.K., Li, Y., Wang, Q.J., Sali, A., Rout, M.P., and Chait, B.T. (2015). A strategy for dissecting the architectures of native macromolecular assemblies. Nat Methods 12, 1135–1138.

Sievers, F., and Higgins, D.G. (2014). Clustal Omega, accurate alignment of very large numbers of sequences. Methods Mol Biol 1079, 105–116.

Sircar, A., Sanni, K.A., Shi, J., and Gray, J.J. (2011). Analysis and modeling of the variable region of camelid single-domain antibodies. J Immunol 186, 6357–6367.

Waterhouse, A.M., Procter, J.B., Martin, D.M., Clamp, M., and Barton, G.J. (2009). Jalview Version 2--a multiple sequence alignment editor and analysis workbench. Bioinformatics 25, 1189–1191.

Webb, B., and Sali, A. (2014). Comparative Protein Structure Modeling Using MODELLER. Curr Protoc Bioinformatics 47, 5 6 1–32.

Wei, X., Decker, J.M., Wang, S., Hui, H., Kappes, J.C., Wu, X., Salazar-Gonzalez, J.F., Salazar, M.G., Kilby, J.M., Saag, M.S., et al. (2003). Antibody neutralization and escape by HIV-1. Nature 422, 307–312.

Wine, Y., Boutz, D.R., Lavinder, J.J., Miklos, A.E., Hughes, R.A., Hoi, K.H., Jung, S.T., Horton, A.P., Murrin, E.M., Ellington, A.D., et al. (2013). Molecular deconvolution of the monoclonal antibodies that comprise the polyclonal serum response. Proc Natl Acad Sci U S A 110, 2993–2998.

Xiang, Y., Shen, Z., and Shi, Y. (2020). Chemical Cross-Linking and Mass Spectrometric Analysis of the Endogenous Yeast Exosome Complexes. Methods Mol Biol 2062, 383–400.

Yu, C., and Huang, L. (2018). Cross-Linking Mass Spectrometry: An Emerging Technology for Interactomics and Structural Biology. Anal Chem 90, 144–165.

Zhu, W., Smith, J.W., and Huang, C.M. (2010). Mass spectrometry-based label-free quantitative proteomics. J Biomed Biotechnol 2010, 840518.

